# Merkel cell polyomavirus protein ALTO modulates TBK1 activity to support persistent infection

**DOI:** 10.1101/2024.04.05.588227

**Authors:** Ranran Wang, Taylor E. Senay, Tiana T. Luo, James M. Regan, Nick Salisbury, Denise Galloway, Jianxin You

## Abstract

While Merkel cell polyomavirus (MCPyV or MCV) is an abundant virus frequently shed from healthy skin, it is one of the most lethal tumor viruses in immunocompromised individuals, highlighting the crucial role of host immunity in controlling MCPyV oncogenic potential. Despite its prevalence, very little is known about how MCPyV interfaces with the host immune response to maintain asymptomatic persistent infection and how inadequate control of MCPyV infection triggers MCC tumorigenesis. In this study, we discovered that the MCPyV protein, known as the Alternative Large Tumor Open Reading Frame (ALTO), effectively primes and activates the STING signaling pathway. It recruits Src kinase into the complex of STING downstream kinase TBK1 to trigger its autophosphorylation, which ultimately activates the subsequent antiviral immune response. Combining single-cell analysis with both loss- and gain-of-function studies of MCPyV infection, we demonstrated that the activity of ALTO leads to a decrease in MCPyV replication. Thus, we have identified ALTO as a crucial viral factor that modulates the STING-TBK1 pathway, creating a negative-feedback loop that limits viral infection and maintains a delicate balance with the host immune system. Our study reveals a novel mechanism by which a tumorigenic virus-encoded protein can link Src function in cell proliferation to the activation of innate immune signaling, thereby controlling viral spread and sustaining persistent infection. Our previous findings suggest that STING also functions as a tumor suppressor in MCPyV-driven oncogenesis. This research provides a foundation for investigating how disruptions in the finely tuned virus-host balance, maintained by STING, could alter the fate of MCPyV infection, potentially encouraging malignancy.

**Author summary:** Merkel cell polyomavirus (MCPyV) is a small DNA virus responsible for the majority cases of Merkel Cell Carcinoma (MCC), a rare yet highly aggressive form of cancer. While MCPyV latently infects a vast majority of individuals, the mechanisms governing its control, persistence, and oncogenic triggers remain obscure. Our research reveals that the MCPyV-derived Alternative Large Tumor Open Reading Frame (ALTO) protein primes and activates the cGAS-STING- TBK1 innate immune pathway within cells. Such activation elicits an antiviral response, effectively curbing MCPyV replication. This phenomenon illustrates the virus’s cunning strategy to exploit cellular mechanisms, ensuring low viral presence while facilitating long-term infection. Consequently, our study sheds light on a novel tactic utilized by MCPyV to maintain a persistent infection in its host.

## Introduction

Merkel cell polyomavirus (MCPyV) has been established as a new human oncogenic virus associated with Merkel cell carcinoma (MCC), a highly aggressive form of skin cancer with a nearly 50% mortality rate [1–5]. The virus carries a ∼5kb circular, double-stranded DNA genome that can be divided into early and late regions [1]. The early region encodes large T (LT), small T (sT), 57kT, and a protein called alternative LT ORF (ALTO) [1, 6–8]. The late region encodes the capsid proteins, VP1 and VP2, and a microRNA [7, 9, 10]. Despite its limited toolkit, MCPyV successfully infects the skin of most humans and establishes long-lasting infections [11, 12]. While MCPyV typically causes asymptomatic infections in the general population [11–14], immunocompromised individuals face a significantly higher risk of developing MCC [14, 15]. In approximately 80% of MCCs, the MCPyV genome is found clonally integrated into the host genome [16, 17]. Integrated MCPyV genomes constitutively express sT and truncated LT (LTT), which function as the major oncogenes that drive MCC tumor growth [2, 17–22]. Therefore, MCPyV integration into the host genome is considered a crucial event in MCPyV-induced oncogenesis [16, 17].

Epidemiological studies have established a strong link between immunosuppression, high MCPyV genome loads, and an increased risk of MCC [14, 23]. These findings highlight the critical role of host immunity in managing MCPyV infection and preventing cancer. In individuals with healthy immune systems, the virus and the human host maintain a delicate balance, allowing persistent infection. However, if the immune system fails to keep this balance, uncontrolled viral replication can occur. This can lead to the integration of the MCPyV genome into the host’s DNA, triggering the development of MCC. However, very little is known about the host innate immune control of MCPyV infection and how dysregulated immune responses may contribute to MCPyV-driven tumorigenesis.

Typical immune responses to viral infection begin with the detection of pathogen-associated molecular patterns (PAMPs) or damage-associated molecular patterns (DAMPs) by pattern recognition receptors (PRRs). The stimulator of interferon genes (STING) is a key mediator of innate immune signaling triggered by PRRs [24, 25]. Upon activation by upstream cytosolic DNA sensors, the endoplasmic reticulum (ER)-resident STING recruits TANK binding kinase 1 (TBK1), which then dimerizes, auto-phosphorylates, and phosphorylates STING. Subsequently, the STING-TBK1 complex translocates through the Golgi complex to the perinuclear lysosomal compartments, where TBK1 phosphorylates interferon regulatory factor 3 (IRF3). Additionally, recruitment of TBK1 by activated STING can also stimulate IκB kinase (IKK) to phosphorylate the NF-κB inhibitor, IκBα, leading to its degradation and the activation of NF-κB. The activated IRF3 and NF-κB then enter the nucleus to induce the transcription of type I interferons (IFNs) and other inflammatory cytokines, thereby establishing an antiviral state [24, 26, 27].

In our previous study, we identified human dermal fibroblasts (HDFs) as the skin cell type which supports productive MCPyV infection and established the first cell culture infection model for the virus [28–31]. Leveraging this system, we probed the early innate immune response to MCPyV. We discovered that MCPyV infection of primary HDFs activates the cGAS-STING-TBK1 pathway, which, in conjunction with NF-κB, activates the expression of IFNs, IFN-stimulated genes (ISGs), and inflammatory cytokines [32, 33]. Our results suggest that the innate immune response initiated by STING signaling is essential for controlling viral infection and facilitating viral persistence. Conversely, we discovered that STING is silenced in MCPyV(+) MCC tumors [34], indicating that the loss of STING function is necessary for MCC tumorigenesis. We propose that the absence of STING function disrupts the MCPyV-host balance, leading to uncontrolled MCPyV replication, which can further promote viral DNA integration and MCC development. Moreover, STING-mediated innate immune responses may serve as an intrinsic barrier to tumorigenesis by linking DNA damage signals, introduced by MCPyV replication, to anti-tumor mechanisms [34, 35]. Therefore, impairing STING function might enable MCPyV-infected cells to bypass these tumor suppressive effects and develop into MCC tumors.

Given that STING is a pivotal component of the antiviral innate immune response [24, 25] and may also act as a tumor suppressor in controlling MCPyV- induced tumorigenesis [32–34], we explored the possibility that MCPyV-encoded proteins could influence STING function. Here we present our findings that identify the MCPyV protein ALTO as the key viral factor that modulates the STING-TBK1 pathway to restrict viral infection activity, thereby preserving a delicate balance with the host’s immune system and enabling persistent infection.

## Results

### ALTO co-operates with STING to induce IFNβ expression

To test whether individual MCPyV encoded-proteins could modulate the STING signaling pathway, we co-transfected HEK293 cells with a STING expression construct and either a construct expressing one of the MCPyV-encoded proteins or an empty vector. A vector expressing RFP was used as a negative control for STING. A highly active STING^R284S^ mutant [36] was used in the first experiment to allow easy detection of the STING downstream signaling. Compared to the RFP control, the STING^R284S^ mutant significantly activated IFNβ mRNA expression (Fig 1A and S1 Fig). More importantly, compared to the empty vector, transfection with the ALTO expression construct dramatically increased IFNβ mRNA expression induced by STING^R284S^ (Fig 1A). In contrast, all other MCPyV-encoded genes showed no effect on STING downstream IFNβ expression (S1 Fig).

**Fig 1.**
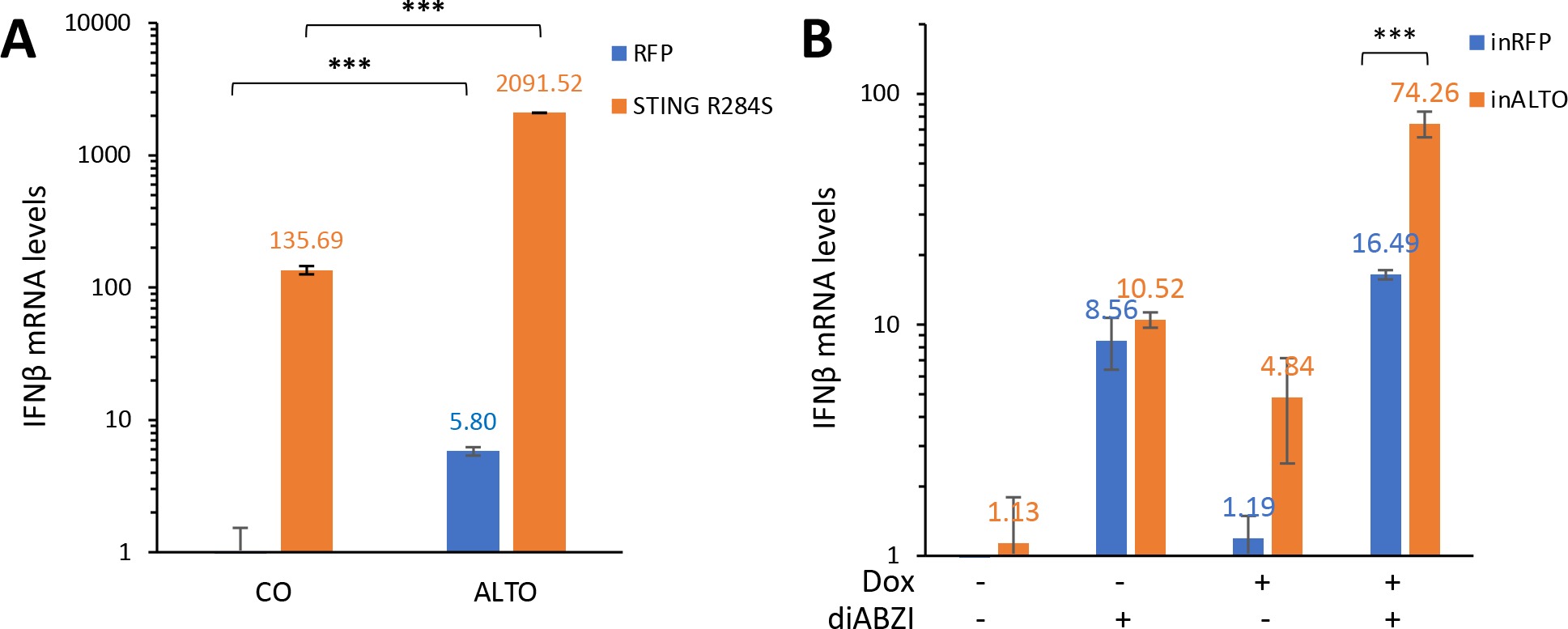
ALTO co-operates with STING to stimulate IFNβ expression. A. HEK293 cells were transfected with plasmids carrying ALTO (or an empty vector control) and STING R284S (or RFP control). Two days post-transfection, mRNA levels of IFNβ was measured using RT-qPCR. One of the values for transfection with vector and RFP was set as 1. **B.** HDF-inRFP and -inALTO cells were mock-treated or induced with Dox and treated with diABZI (or DMSO control) for 19 hours. mRNA levels were measured using RT-qPCR. The value from HDF-inRFP cells mock- and DMSO-treated was set as 1. Error bars indicate standard deviation from three independent samples. ***p<0.001.

We then tested how ALTO expression impacts the STING pathway in primary HDFs that support MCPyV infection [28]. We generated stable HDF cells to allow inducible expression of either RFP (inRFP) or ALTO (inALTO) by Doxycycline (Dox). Dox treatment was then used to induce RFP or ALTO expression in these cells (Fig 1B). In MCPyV infected cells, ALTO is likely to function in the presence of replicated viral DNA, which could stimulate the STING-TBK1 pathway. Therefore, we sought to simulate this biological setting by treating HDF-inRFP or -inALTO cells with and without Dox in the absence and presence of the STING agonist diamidobenzimidazole (diABZI). In the HDF- inRFP cells, diABZI treatment could stimulate IFNβ expression, but RFP expression does not have additional effect (Fig 1B). However, in HDF-inALTO cells, while both diABZI treatment and ALTO induction were found to moderately up-regulate IFNβ mRNA expression, combining ALTO induction with diABZi treatment resulted in a significant increase in IFNβ expression (Fig 1B).

Together, our studies showed that ALTO can stimulate STING downstream signaling.

### MCPyV ALTO stimulates TBK1 autophosphorylation and STING stabilization

We next tested if ALTO stimulates IFNβ expression by altering the status of STING and its downstream components including TBK1, IRF3, and NF-κB. HDF-inRFP or -inALTO cells were treated with diABZI as a positive control to show how STING activation changed the phosphorylation status of these factors (Fig 2A). As expected, in HDF-inRFP and -inALTO cells, diABZI treatment led to a substantial activation of STING and downstream signaling as indicated by TBK1^S172^ autophosphorylation and phosphorylation of STING^S366^, IRF3^S386^, and the NF-κB subunit p65^S536^ (Fig 2A, lanes 2 and 6). STING phosphorylation is required for its trafficking and subsequent ubiquitination and degradation [37, 38]. Consistently, we observed a sharp reduction of STING protein level in diABZI-treated cells (Fig 2A, left panel, lanes 2 and 6). However, in HDF-inALTO cells treated with only Dox but not with diABZI, we detected an increased level of STING but not the phosphorylated STING^S366^ (Fig 2A, compare lane 7 to lanes 1, 3, and 5). Immunofluorescent (IF) staining confirmed that, compared to HDF- inRFP cells or mock-induced HDF-inALTO cells, more STING was accumulated in Dox-induced HDF-inALTO cells both in the presence and absence of diABZI (Fig 2B). More strikingly, TBK1^S172^ phosphorylation was markedly elevated by ALTO expression even in the absence of diABZI (Fig 2A, left panel, compare lane 7 to lanes 1, 3, and 5). TBK1^S172^ phosphorylation was not detected in HDF- inRFP cells treated with Dox (Fig 2A, left panel, lane 3), suggesting that ALTO expression was responsible for inducing TBK1^S172^ phosphorylation. Dual treatment of HDF-inALTO cells with Dox and diABZI did not significantly enhance TBK1^S172^ autophosphorylation over Dox treatment alone (Fig 2A, left panel, compare lane 7 and 8). The level of total TBK1 protein was reduced in the HDF- inRFP cells treated with diABZI (Fig 2A, left panel, compare lanes 2 and 4 with lanes 1 and 3), consistent with normal degradation following activation [39]. In HDF-inALTO cells, the level of TBK1 protein was further reduced by ALTO expression, both with and without diABZI treatment (Fig 2A, left panel, compare lanes 7-8 to lanes 2, 4, and 6). TBK1^S172^ autophosphorylation is needed for suppressor of cytokine signaling 3 (SOCS3)-mediated proteasomal degradation [39]. Therefore, it is likely that ALTO-induced TBK1^S172^ autophosphorylation stimulated its degradation, thereby reducing the amount of TBK1 available for STING phosphorylation. Because STING phosphorylation triggers its degradation [37, 38], the lack of STING phosphorylation could inhibit its degradation and cause the observed accumulation of STING in the ALTO-expressing cells.

**Fig 2.**
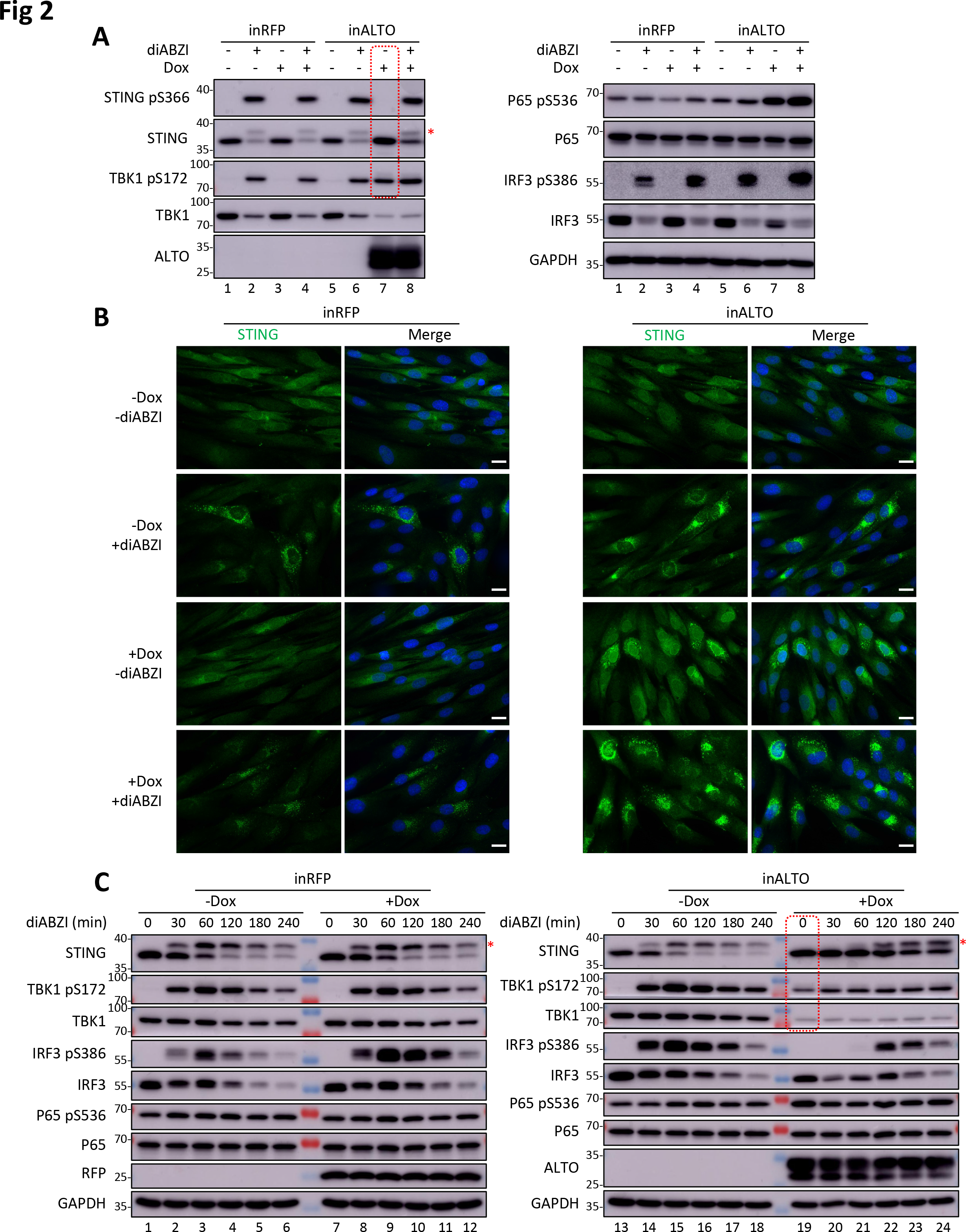
ALTO stimulates TBK1^S172^ autophosphorylation and STING stabilization. A. HDF-inALTO or -inRFP cells were mock-treated or induced with Dox for 24 hours, then stimulated with diABZI (or DMSO control) for 16 hours. Whole cell lysates were resolved by SDS/PAGE and immunoblotted with the indicated antibodies. **B.** HDF-inALTO or - inRFP cells were mock-treated or induced with Dox for 7 hours, then stimulated with diABZI (or DMSO control) for an additional 19 hours. Cells were fixed and immune-stained for STING, and counterstained with DAPI. Scale bar, 20 µm. **C.** HDF-inALTO or - inRFP were mock-treated or induced with Dox for 40 hours, then stimulated with diABZI (or DMSO control) for the indicated times. Whole cell lysates were resolved by SDS/PAGE and immunoblotted with indicated antibodies. Red asterisks indicate phosphorylated STING.

Consistent with a recent study [8], we also found that ALTO expression slightly enhanced p65^S536^ phosphorylation (Fig 2A, right panel).

To mimic the biological setting in which ALTO function might be affected by the dynamics of MCPyV infection, we treated HDF-inRFP or -inALTO cells with and without Dox in both the absence and presence of diABZI for varying amounts of time. In both types of cells not treated with Dox, diABZI stimulated the phosphorylation of STING, TBK1^S172^, and IRF3^S386^, and to a lesser degree, p65^S536^ phosphorylation (Fig 2C). In HDF-inALTO cells, Dox-induced ALTO expression in the absence of diABZI again led to a significantly increased TBK1^S172^ phosphorylation, reduced TBK1 protein level, and stabilization of STING (Fig 2C, compare lane 19 to lane 13). During the earlier time points (0, 30, 60 min) of diABZI treatment of Dox-induced HDF-inALTO cells, there was a pronounced increase in STING protein level, but a reduction in phosphorylated STING compared to HDF-inRFP cells or HDF-inALTO cells not treated with Dox (Fig 2C, Compare lanes 19-21 to lanes 1-3, 7-9, and 13-15). This is consistent with our observation that ALTO can stabilize STING but does not stimulate its phosphorylation (Fig 2A). As discussed above, this observation could be attributed to the fact that ALTO expression greatly reduced the level of TBK1 needed for STING phosphorylation (Fig 2C, compare lanes 19-24 to lanes 13- 18). Interestingly, after Dox-induced HDF-inALTO cells were treated with diABZI for longer periods (120, 180, 240 min), phosphorylated STING levels substantially increased (Fig 2C, compare lanes 22-24 to lanes 4-6, 10-12, and 16-18). It is possible that at these later time points, STING accumulated in these cells was eventually phosphorylated by either the residual TBK1 present in the cells or alternative kinases.

We also found that even a trace amount of ALTO expressed for a very brief period (4h) could lead to robust TBK1^S172^ autophosphorylation and STING accumulation (S2 and S3 Fig), suggesting that this ALTO function may play a physiologically important role in MCPyV infected cells, which was confirmed in our studies described below. TBK1 protein level dropped strikingly after extended ALTO induction and TBK1^S172^ autophosphorylation (S3 Fig), supporting the role of TBK1^S172^ autophosphorylation in stimulating its ubiquitination and degradation [39]. Together, our studies suggest that ALTO first induces TBK1 autophosphorylation and reduction, subsequently causing STING accumulation and activation.

### ALTO forms a complex with STING and TBK1

To understand how ALTO could so effectively stimulate TBK1 autophosphorylation and STING stabilization, we decided to test if this viral protein interacts with STING and TBK1. We first performed ALTO Co-IP using HDF-inALTO cells induced with Dox. While STING was not clearly Co-IPed with ALTO, both TBK1 and phosphorylated TBK1^S172^ were found to be efficiently pulled down with ALTO antibody (Fig 3A). C-terminal HA- and FLAG-tagged ALTO could also efficiently Co-IP phosphorylated TBK1^S172^ (S4 Fig).

**Fig 3.**
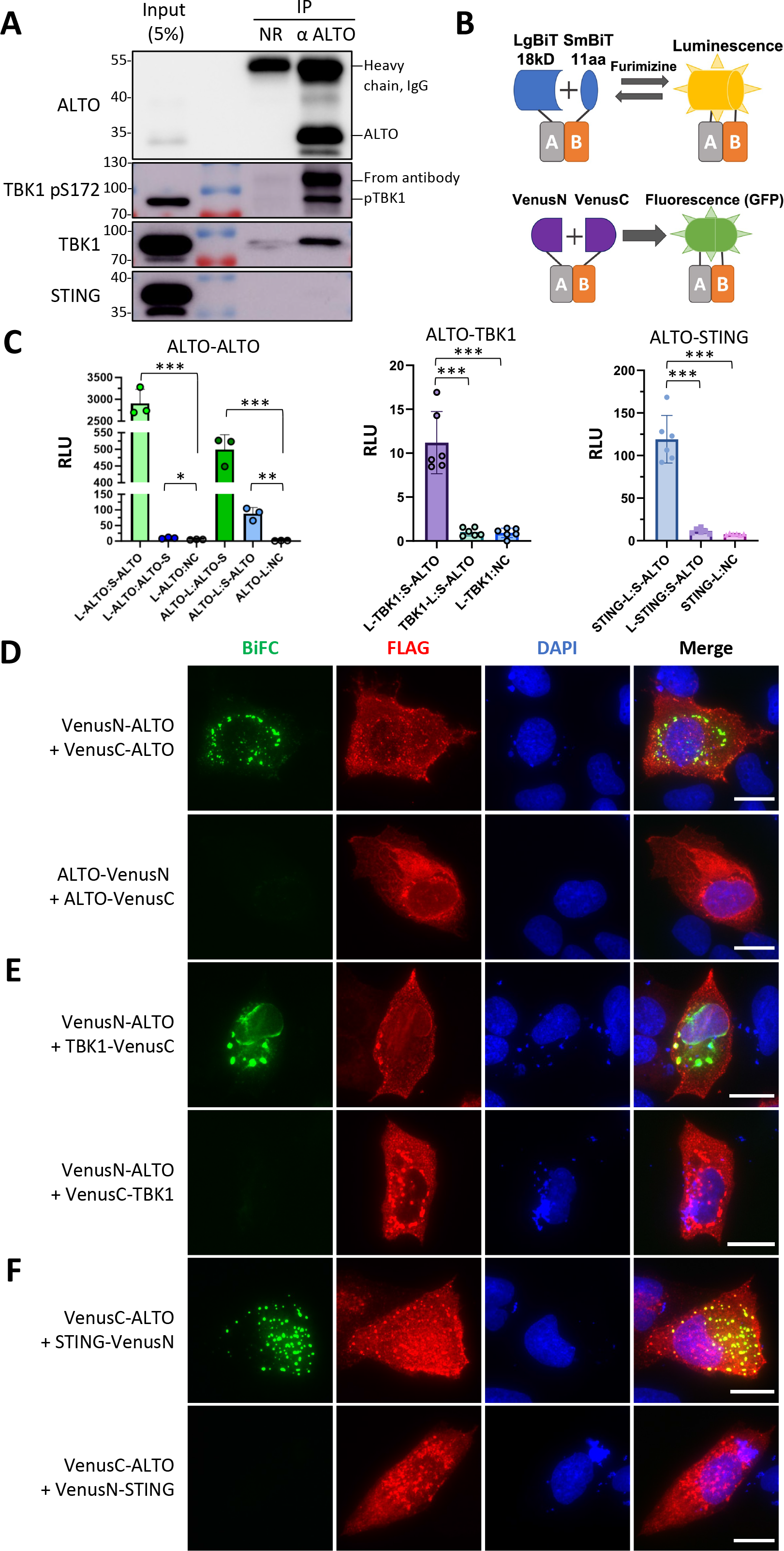
ALTO interacts with STING and TBK1. A. HDF-inALTO cells were induced with 0.01μg/mL Dox for 40 hours. Whole cell lysates were incubated with normal rabbit IgG or anti-ALTO antibody and immunoprecipitated with Protein G agarose. Co- immunoprecipitants were resolved by SDS/PAGE and immunoblotted with the indicated antibodies. **B.** Schematic diagram depicting the NanoBiT and BiFC structural complementation systems. NanoBiT reversibly reconstitutes NanoLuciferase and produces luminescence in live cells, whereas BiFC irreversibly reconstitutes Venus to produce GFP fluorescence. **C.** HEK293 cells were transfected with pairs of plasmids carrying NanoBiT-tagged ALTO, STING, and TBK1 or the Small BiT negative control construct. At 20 hours (ALTO-TBK1) or 24 hours (ALTO-ALTO and ALTO-STING) post- transfection, NanoGlo reagent was added, and luminescence was measured after a 15- minute (ALTO-ALTO) or 2-hour (ALTO-STING and ALTO-TBK1) incubation. Points indicate replicate wells from a single experiment, bars indicate means, and error bars indicate standard deviations. L-protein and protein-S (and similar) indicate the named protein tagged with the LgBiT at its N-terminus or SmBiT at its C-terminus, respectively. ***p<0.001; **p<0.01; *p<0.05. **D.** U2OS cells were transfected with pairs of plasmids carrying FLAG-tagged Venus (N-terminal half or C-terminal half) fused ALTO constructs. At 48 hours post-transfection, cells were fixed and immunostained for FLAG, and counterstained with DAPI. Scale bar, 20 μm. **E.** U2OS cells were transfected with pairs of plasmids carrying FLAG-tagged Venus (N-terminal half or C-terminal half) fused ALTO and TBK1, respectively. At 36 hours post-transfection, cells were fixed and immunostained for FLAG, and counterstained with DAPI. **F.** U2OS cells were transfected with pairs of plasmids carrying FLAG-tagged Venus (N-terminal half or C- terminal half) fused ALTO and STING, respectively. At 24 hours post-transfection, cells were fixed and immunostained for FLAG, and counterstained with DAPI. Scale bar, 20 μm.

We then applied two structural complementation-based cellular approaches to examine the interaction between ALTO, STING and TBK1: NanoLuc Binary Technology (NanoBiT) and bimolecular fluorescence complementation (BiFC) (Fig 3B). The NanoBiT system is a split luciferase- based reporter assay designed to sensitively detect protein-protein interactions in live cells [40]. Proteins of interest are tagged with a piece of Nano Luciferase, called Large (LgBiT) or Small BiT (SmBiT), positioned at the N- or C-terminus.

When LgBiT and SmBiT are brought into close proximity by a protein-protein interaction, the Nano Luciferase is reversibly reconstituted and will produce luminescence in the presence of the cell-permeable substrate furimazine (Fig 3B). To assess ALTO’s interaction with STING and TBK1 in this system, we generated fusions with all possible combinations of BiT size and location for all proteins, for a total of four constructs per protein. All combinations of LgBiT and SmBiT pairs were screened for all interacting pairs. Interestingly, we found that ALTO pairs tagged on the same terminus (i.e., LgBiT-ALTO:SmBiT-ALTO, and ALTO-LgBiT:ALTO-SmBiT) produced relative luminescence readings several hundred times higher than negative interaction controls (Fig 3C). These data demonstrate that ALTO strongly interacts with itself in living cells. The two opposite-terminal pairs, ALTO-LgBiT:SmBiT-ALTO and LgBiT-ALTO:ALTO- SmBiT also luminesced significantly more than their LgBiT negative control; however, the overall magnitude of the luminescence was much lower than the same-terminal pairs, and C-terminal tagged ALTO was apparently expressed at a notably lower level than N-terminal tagged ALTO (S5A Fig). Since ALTO has a predicted C-terminal transmembrane domain, this result indicates that membrane localization and orientation may impact ALTO’s oligomerization. SmBiT-ALTO was also found to positively interact with both LgBiT-TBK1 and STING-LgBiT at levels significantly above both negative controls and opposite-terminal controls (Fig 3C). This interaction is considered genuine, as the luminescence readings for SmBiT-ALTO:LgBiT-TBK1 and SmBiT-ALTO:STING-LgBiT were more than tenfold higher than the average from the LgBiT-tagged construct co-transfected with the SmBiT negative control construct, thereby meeting the manufacturer’s criteria for a confirmed protein-protein interaction and demonstrating the presence and specificity of these interactions. NanoBiT-fused TBK1, but also to a lesser degree NanoBiT-fused STING, were not expressed as efficiently as the ALTO fusion proteins, which likely explains the overall lower relative luminescence in these assays (S5B and S5C Fig). Taken together, these data show that, in addition to oligomerizing, ALTO is in close proximity to TBK1 and STING in cells and is therefore likely to form a complex with these host proteins.

The ALTO-ALTO NanoBiT interaction signal is significantly higher than those for ALTO-TBK1 and ALTO-STING. This is likely because ALTO tends to self-oligomerize and co-localize within cells (see below), leading to a constant proximity of the tagged proteins and a higher likelihood of reconstituting the luciferase enzyme. In contrast, interactions between ALTO and TBK1 or STING are more transient and less frequent due to the dynamic nature of their interactions and the potential for ALTO to reduce TBK1 levels. Additionally, the untagged endogenous TBK1 and STING might preferentially interact with ALTO, thereby reducing the luminescence signals for ALTO-TBK1 and ALTO-STING NanoBiT. Based on these observations, we decided to further investigate the ALTO-ALTO, ALTO-STING, and ALTO-TBK1 interactions using a complementary approach, Bi-FC. This method renders the interaction between the proteins irreversible, thus making it more easily observable.

In a BiFC assay, two interacting protein partners are fused individually to complementary nonfluorescent fragments of a fluorescent reporter Venus and expressed in live cells [41] (Fig 3B). Interaction of these two proteins will bring the nonfluorescent fragments within close proximity, allowing the Venus protein to reform its native three-dimensional structure and emit fluorescent signal [42]. Therefore, this technique enables direct visualization of protein interactions within the cell through the intensity and distribution of the fluorescent signal [41]. To test the ALTO-ALTO, ALTO-STING, and ALTO-TBK1 protein interactions, these molecules were fused individually to either the N-terminal portion (VenusN) or C-terminal portion (VenusC) of the divided Venus protein. Coexpression of constructs encoding ALTO with N-terminal tagged VenusN and VenusC generated a bright BiFC signal, which is absent in cells co-transfected with constructs encoding ALTO with VenusN and VenusC fused to its C-terminus (Fig 3D, S6 and S7 Fig). This result provides additional evidence to demonstrate that ALTO can oligomerize in cells. Co-expression of VenusN-ALTO with TBK1-VenusC or VenusC-ALTO with STING-VenusN also generated a strong BiFC positive signal (Fig 3E and 3F). Switching the VenusC tag from the C to N-terminus of TBK1 or the VenusN tag from the C to N-terminus of STING abolished the BiFC signal (Fig 3E and 3F). Since the differently tagged ALTO, STING and TBK1 were expressed equally well in cells (S7-S9 Fig), they served as excellent negative controls to support that the BiFC signals we observed likely resulted from true ALTO-ALTO, ALTO-TBK1 and ALTO-STING interaction, and not from overexpression of the proteins tested (Fig 3D, 3E, 3F, S7-S9 Fig).

### ALTO recruits Src to the TBK1 complex to stimulate its autophosphorylation

To better understand how ALTO could stimulate TBK1 autophosphorylation, we examined the ALTO protein sequence and identified 13 proline-X-X-proline (PXXP) motifs (Fig 4A). Interestingly, the STING sequence also contains one PXXP motif, which has been shown to bind to the SH3 domain of the tyrosine kinase Src [43]. This binding recruits the kinase to phosphorylate TBK1^Y179^, stimulating TBK1 autophosphorylation and activation [43]. Since the 13 PXXP motifs present in ALTO are mostly located on the disordered surface of the ALTO structure as predicted by AlphaFold2 [44] (Fig 4A), we tested whether ALTO could interact with Src. C-terminal HA- and FLAG-tagged ALTO was able to pull down a small amount of Src (S4 Fig). In the reciprocal Co-IP experiment, Src antibody Co-IP was performed using HDFs inducibly expressing ALTO (or RFP) either mock-treated or stimulated with diABZI (Fig 4B and S10 Fig). Both mouse and rabbit Src antibodies successfully precipitated ALTO, but this occurred only in cells treated with diABZI (Fig 4B and S10 Fig), suggesting that specific stimuli, such as replicating MCPyV DNA, might be necessary to facilitate the interaction between Src and ALTO. Upon treatment with diABZI, Src antibodies were able to Co-IP TBK1, particularly its S172 phosphorylated form (Fig 4B and S10 Fig). We suspect that Src interacts with both ALTO and TBK1, even in the absence of diABZI treatment (S4 Fig). However, the introduction of upstream stimuli may bolster this interaction or stabilize a transient association that might not otherwise endure the stringent conditions of the co-IP process. More importantly, compared to the cells expressing RFP, the amount of TBK1 and phosphorylated TBK1^S172^ complexed with Src was substantially increased in the cells expressing ALTO (Fig 4B and S10 Fig). Considering the observation that ALTO interacts with TBK1 (Fig 3 and S4 Fig), this result suggests that ALTO may utilize its multiple PXXP motifs to recruit Src into the TBK1 complex.

**Fig 4.**
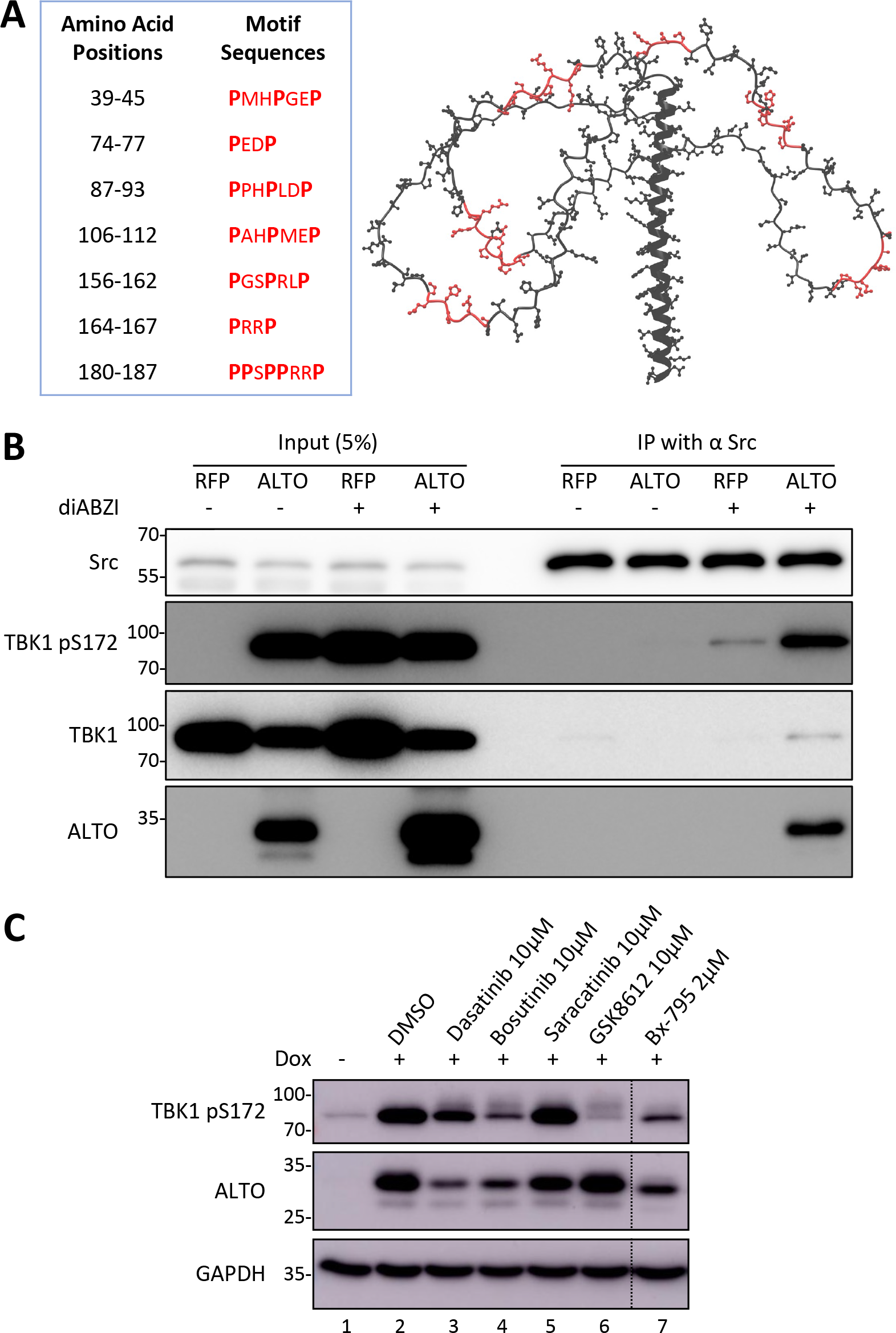
ALTO recruits Src into complex with TBK1 to stimulate its autophosphorylation. A. AlphaFold2 model of ALTO protein, with the Src-binding P-X-X-P motifs denoted in red. The table provides details for P-X-X-P motif sequences and positions. **B.** HDF-inALTO or -inRFP cells were mock-treated or induced with Dox for 10.5 hours, then stimulated with diABZI (or DMSO control) for 1.5 hours. Whole cell lysates were incubated with mouse anti-Src antibody and immunoprecipitated with Protein G agarose. Immunoprecipitants were resolved by SDS/PAGE and immunoblotted with the indicated antibodies. **C.** HDF-inALTO cells were mock-treated or pre-treated with the indicated doses of Src/SFK or TBK1 small molecule inhibitors for 1 hour, then mock-treated or induced with 0.1 μg/mL Dox for 4 hours. Whole cell lysates were resolved by SDS/PAGE and immunoblotted with the indicated antibodies.

Given that Src has been demonstrated to prime TBK1 for autophosphorylation and activation [43], the capacity of ALTO to facilitate the formation of the Src-TBK1 complex may elucidate ALTO’s effectiveness in stimulating TBK1 autophosphorylation (Fig 2, S2 and S3 Fig). To test if Src plays a key role in ALTO-induced TBK1^S172^ autophosphorylation, we treated HDFs inducibly expressing ALTO with Src family kinase (SFK) inhibitors as well as TBK1 inhibitors (as positive control) (Fig 4C). Cell viability tests showed that all three Src/SFK inhibitors were well tolerated by the HDF-inALTO cells (S11 Fig). The TBK1 inhibitors GSK8612 and Bx-795 could both block TBK1^S172^ autophosphorylation (Fig 4C). The Src/SFK inhibitors Dasatinib and Bosutinib also significantly suppressed TBK1^S172^ autophosphorylation, whereas Saracatanib showed very little effect (Fig 4C). The Src/SFK inhibitors did not reduce TBK1 autophosphorylation as effectively as the TBK1 inhibitors, likely because in this experiment we focused on monitoring TBK1^S172^ autophosphorylation, which is downstream of the Src-mediated TBK1^Y179^ phosphorylation [43]. Nevertheless, our inhibitor data suggest that Src plays a crucial role in ALTO-induced TBK1^S172^ autophosphorylation. This supports the model where ALTO facilitates the recruitment of Src into the complex with TBK1, thereby stimulating its autophosphorylation and activation.

### The induction of IFN signaling by MCPyV infection may be attributed to the stabilization of STING

We next started to examine the functional impact of ALTO-induced TBK1 autophosphorylation and STING stimulation during MCPyV infection. As established in our previous studies [32, 33], infecting HDFs with a moderate dose of MCPyV (10^8^ viral genome equivalents of MCPyV virions per 96-well) triggered a robust expression of IFNβ and key ISGs such as Viperin, Oas1, ISG54, and Mx1 (Fig 5A). The IFN signaling induced by MCPyV is significantly more robust than the signaling triggered by diABZI (Fig 5A). Surprisingly, co-treating cells with MCPyV and diABZI did not enhance, but instead dramatically suppressed, the IFN signaling induced by MCPyV (Fig 5A). Given that IFN stimulation was significantly higher, yet STING mRNA level was lower in MCPyV-treated cells compared to those treated with diABZI (Fig 5A), we decided to examine STING and its downstream signaling molecules at the protein level. Both diABZI and MCPyV treatments stimulated phosphorylation of TBK1^S172^ and IRF3^S386^, with MCPyV treatment additionally inducing p65^S536^ phosphorylation (Fig 5B). The STING protein level was reduced by diABZI, which is known to stimulate STING phosphorylation and subsequent degradation [45, 46]. While MCPyV infection activates TBK1 stronger than diABZI treatment, it downregulates STING less (Fig 5B and 5C), consistent with our previous findings that ALTO stabilizes STING.

**Fig 5.**
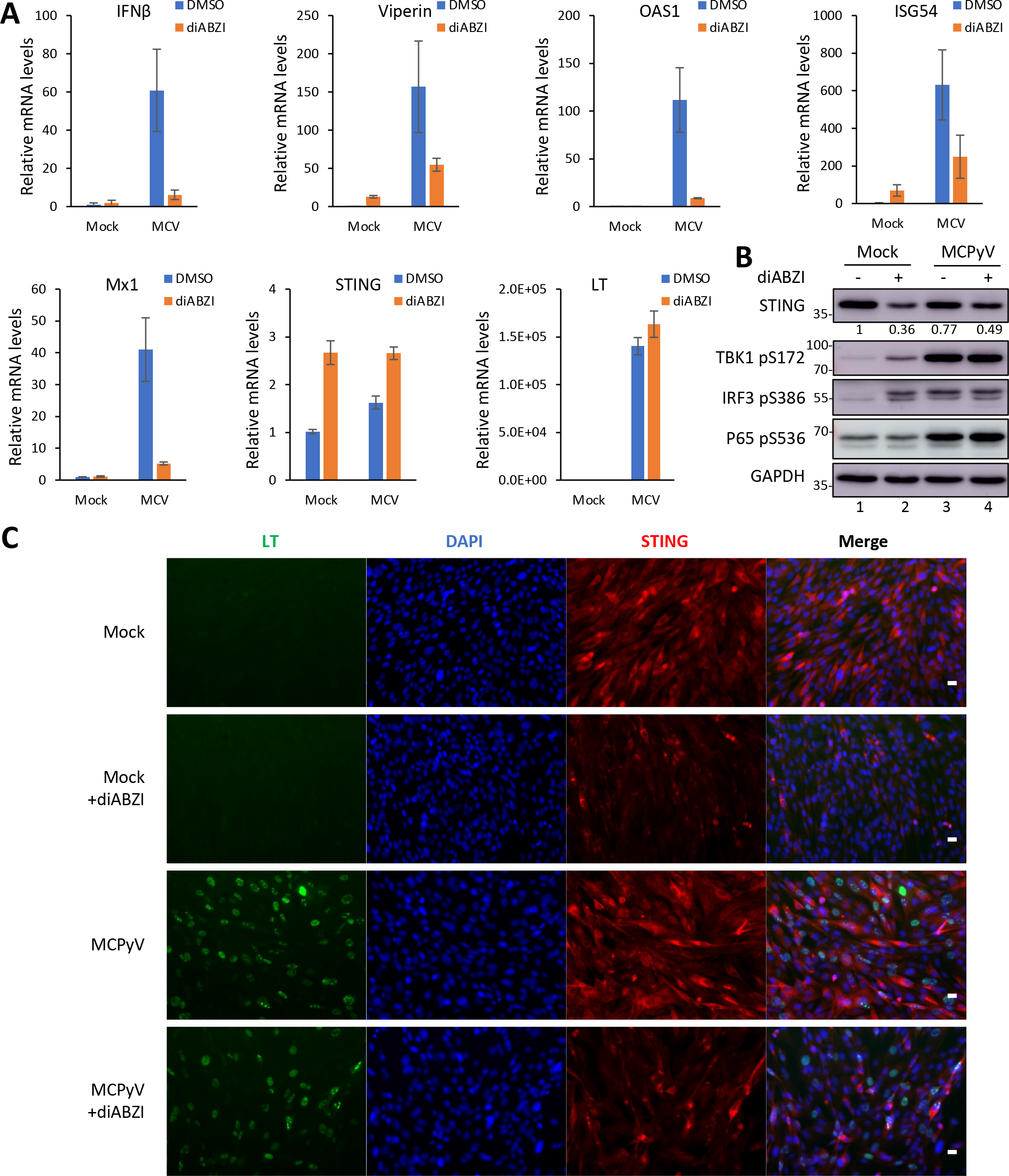
MCPyV rescues STING levels to stimulate IFN signaling. A. Normal HDFs mock-infected or infected with MCPyV were stimulated with diABZI (or DMSO control) at the time FBS was added (2 days post-infection). At 5 days post-infection, cells were harvested, and mRNA levels were quantified by RT-qPCR. The value for mock-infected and DMSO-treated HDFs was set as 1. Error bars indicate standard deviation from 3 replicates. **B.** Normal HDFs were mock-infected or infected with MCPyV, then stimulated with diABZI (or DMSO control) at the time FBS was added (2 days post-infection). Whole cell lysates were collected at 5 days post-infection, resolved by SDS/PAGE, and immunoblotted with the indicated antibodies. **C.** Normal HDFs were mock-infected or infected with MCPyV, then stimulated with diABZI (or DMSO control) at the time FBS was added (2 days post-infection). Cells were fixed at 5 days post-infection and immunostained for LT and STING, and counterstained with DAPI. Scale bar, 20 μm.

This result suggests that the virus may stabilize the STING protein. Compared to cells treated with diABZI, those treated with MCPyV transcribed less STING mRNA but maintained a higher level of STING protein (Fig 5A, 5B and 5C), again supporting the virus’s ability to stabilize STING protein. In the cells treated with both MCPyV and diABZI, MCPyV infection was able to partially counteract the STING degradation induced by diABZI (Fig 5B and 5C). Collectively, these studies suggest that MCPyV infection both activates and stabilizes STING. In light of our previous experiments, these findings suggests that the effect of MCPyV might be attributed to the role of ALTO in stabilizing STING.

### ALTO inhibits MCPyV infection activity

To best understand the potential magnitude and timing of ALTO’s effect during MCPyV infection, we analyzed changes in ALTO protein levels throughout the course of MCPyV infection in the HDF model. ALTO expression became detectable 4-6 days post-infection, peaking on day 5 (Fig 6A). Subsequently, we generated an ALTO null (ALTOnull) MCPyV variant to assess ALTO’s impact on MCPyV infectivity. In HDFs infected with varying titers of wild type (WT) or ALTOnull MCPyV, the ALTOnull virus consistently showed enhanced replication (30-60% better) compared to WT MCPyV (Fig 6B). LT and VP1 were produced at similar levels by WT and ALTOnull MCPyV, indicating that the ALTOnull mutation does not affect the expression of LT and VP1 during infection (S12A Fig). These data suggest that the ALTO protein encoded by the WT virus might suppress viral replication activity. To further investigate this function of ALTO, we infected HDF-inALTO cells with ALTOnull MCPyV. We observed that inducing ALTO in these cells could suppress viral infection by 55-75% (Fig 6C). Both western blotting and IF analysis of HDF-inALTO cells revealed that while Dox treatment increased ALTO expression, the level of MCPyV LT significantly decreased in cells treated with Dox (Fig 6D and S12B Fig). These findings confirm that exogenous ALTO, driven by an inducible promoter, can inhibit MCPyV replication activity.

**Fig 6.**
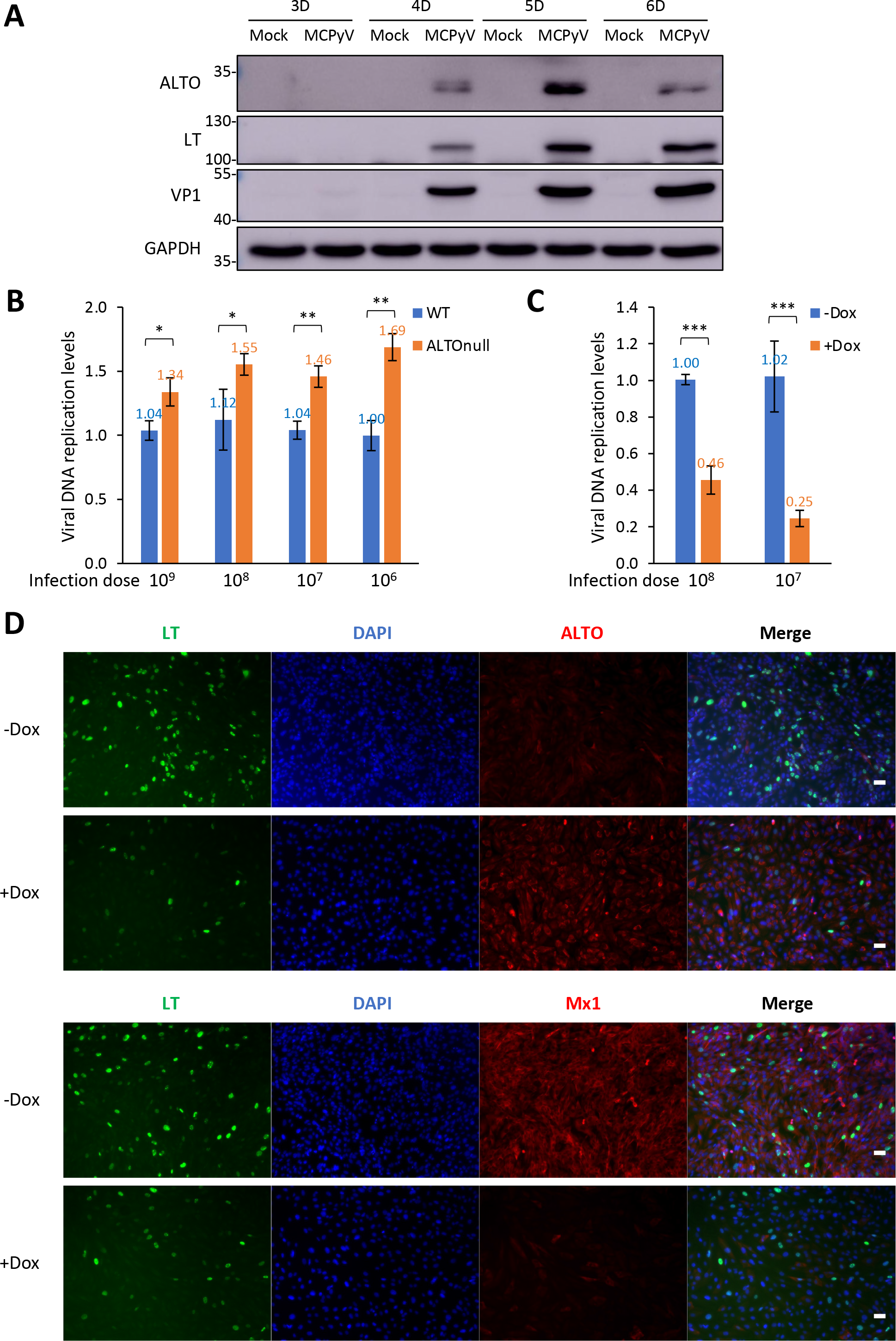
MCPyV ALTO suppresses viral infection activity. A. Normal HDFs were mock- infected or infected with 6x10^8^ copies of MCPyV per 48-well. Whole cell lysates were collected at the indicated timepoints post-infection. Lysates were resolved by SDS/PAGE and immunoblotted with the indicated antibodies. **B.** Normal HDFs were infected with the indicated titers of WT or ALTOnull MCPyV per 96-well. Cells were harvested and DNA extracted at 5 days post-infection. Viral replication was quantified by qPCR using the NCRR-specific primers and normalized to genomic GAPDH levels. Bars indicate mean; error bars indicate standard deviation from 3 replicates. **p<0.01; *p<0.05. **C.** HDF-inALTO cells were infected with the indicated titers of ALTOnull MCPyV per 96-well. Cells were mock-treated or induced with Dox at 2- and 4-days post- infection. Cells were harvested and DNA extracted at 5 days post-infection. Viral replication was quantified by qPCR using the NCRR specific primers and normalized to genomic GAPDH levels. Bars indicate mean; error bars indicate standard deviation from 3 replicates. ***p<0.001. **D.** HDF-inALTO cells were infected with ALTOnull MCPyV. Cells were mock-treated or induced with Dox at 2- and 4-days post-infection. Cells were fixed on day 5 post-infection and stained for LT and either ALTO or Mx1, and counterstained with DAPI. Scale bar, 50 μm.

As indicated by Mx1 IF staining, the reduced MCPyV infection subsequently impaired IFNβ downstream ISG expression (Fig 6D). However, no significant difference was detected in virus-induced phosphorylation of STING^S366^, TBK1^S172^, IRF3^S386^, or p65^S536^ in HDFs infected with either WT or ALTOnull MCPyV (S12 Fig). Given that the Western blotting analysis was conducted on a mixed population of both infected and uninfected cells, we reasoned that examining individual cells could allow for a more accurate profiling of ALTO’s impact on the STING-TBK1 pathway during viral infection. IF staining of WT MCPyV-infected HDFs revealed that cells exhibiting a positive ALTO signal often displayed weaker LT signal (Fig 7A). Additionally, we employed ImmunoFISH to double-stain WT MCPyV-infected HDFs with an ALTO antibody and MCPyV DNA probes [28](Fig 7B). Compared to ALTO-negative cells, many ALTO-positive cells exhibited a reduced FISH signal for the replicated MCPyV genome (Fig 7B and S13 Fig). In contrast, ALTOnull MCPyV-infected cells displayed a stronger MCPyV FISH signal than those infected with WT MCPyV (Fig 7B). Quantification of the ImmunoFISH signal in individual cells further confirmed that ALTO-positive HDFs infected with WT MCPyV exhibited a significantly lower viral replication FISH signal –by ∼50%– compared to ALTO- negative HDFs, regardless of infection with WT or ALTOnull MCPyV (Fig 7C). This result underscores ALTO’s role in repressing MCPyV infection at the single cell level (see discussion).

**Fig 7.**
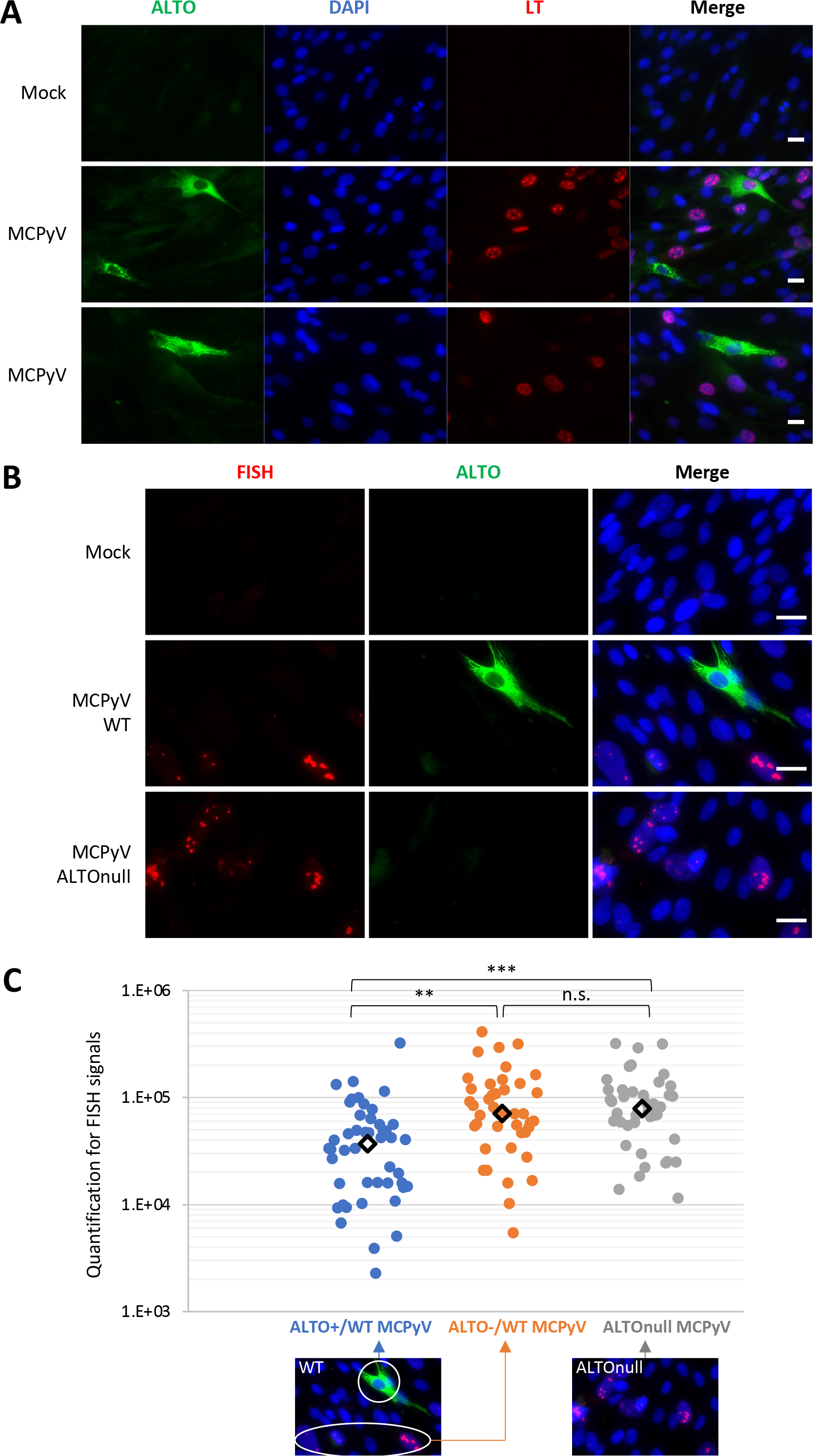
ALTO suppresses MCPyV infection at the single-cell level. A. Normal HDFs were either mock-infected or infected with 6x108 viral genome equivalents of MCPyV virions per well in a 48-well plate. Cells were fixed on day 4 post-infection, immunostained for ALTO and LT, and counterstained with DAPI. Scale bar: 20 μm. **B.** Normal HDFs were either mock-infected or infected with 6x108 viral genome equivalents of WT or ALTOnull MCPyV virions per well in a 48-well plate. Cells were fixed on day 5 post-infection, immunostained for ALTO, subjected to FISH using a MCPyV probe, and counterstained with DAPI. Scale bar: 20 μm. **C.** The FISH signals within individual cells treated as in B were quantified using ImageJ. Shown are the quantification results for the FISH signal in individual ALTO-positive and ALTO-negative HDFs infected with WT MCPyV, as well as HDFs infected with ALTOnull MCPyV. Each type of cell is indicated with color-coded arrows. More than 40 cells were analyzed for each sample group to achieve a statistically meaningful sample size. Central diamonds represent the median values of the data groups. ***p<0.001; **p<0.01; n.s. = not significant.

### TBK1 inhibition stimulates MCPyV replication

Given the potent ability of ALTO to stimulate TBK1^S172^ autophosphorylation (Fig 2, S2 and S3 Fig), we investigated the effects of blocking TBK1 autophosphorylation on MCPyV infection and the role of ALTO in this process. Treating HDF stable cells that can inducibly express ALTO with the TBK1 inhibitor GSK8612 abrogated ALTO-induced TBK1^S172^ autophosphorylation in a dose-dependent manner (Fig 8A, compare lanes 3-6 to lane 2). Compared to the mock- treated control, Dox induction of ALTO in HDF-inALTO cells significantly repressed ALTOnull MCPyV infective activity (Fig 8B). Treatment with GSK8612 alone markedly enhanced viral infectivity (by up to 51%), demonstrating that inhibiting the TBK1 downstream innate sensing pathway can enhance MCPyV infection and spread (Fig 8B). In cells with Dox-induced ALTO expression, treatment with GSK8612 further rescued an additional 23% of the viral replication compared to cells without ALTO induction (Fig 8B). These results highlight TBK1’s critical role in mediating ALTO’s function in controlling MCPyV infection.

**Fig 8.**
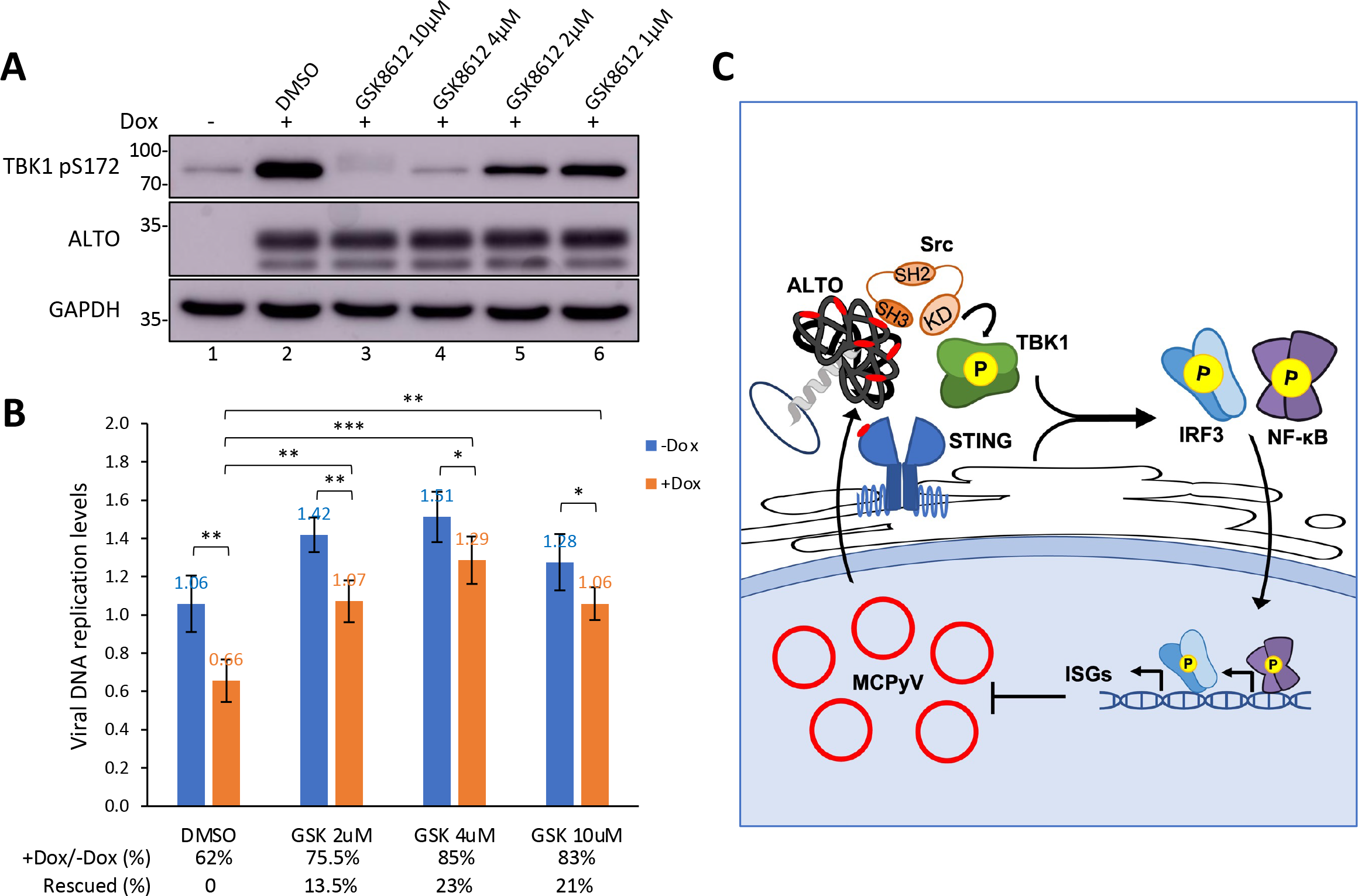
TBK1 inhibition reactivates MCPyV infection activity. A. HDF-inALTO cells were treated with the indicated Src/SFK or TBK1 inhibitors (or DMSO control) for one day, then treated with 0.05 μg/mL Dox for two additional days. Whole cell lysates were resolved by SDS/PAGE and immunoblotted with the indicated antibodies. **B.** HDF- inALTO cells were infected with ALTOnull MCPyV. Beginning 2 days post-infection, the cells were treated with the TBK1 inhibitor GSK8612 (or DMSO control) at the indicated doses, then treated with 0.05 μg/mL Dox at 3 days post-infection. At 5 days post-infection, cells were harvested and DNA extracted. Viral replication was quantified by qPCR using the NCRR-specific primers and normalized to genomic GAPDH. The value from one group of uninduced cells treated with DMSO was set as 1. Bars indicate mean from three replicates; error bars indicate standard deviation. Percentages reflect relative replication in Dox-induced cells compared to their respective uninduced controls. ***p<0.001; **p<0.01; *p<0.05. **C.** ALTO stimulates STING-TBK1 signaling to negatively regulate MCPyV infection. The MCPyV protein ALTO recruits Src via its PXXP motifs (red) to phosphorylate TBK1, leading to STING stabilization and activation. Activated STING induces an enhanced downstream ISG response via IRF3 and NF-κB. This antiviral response provides a negative feedback mechanism to control MCPyV propagation and persistence.

## Discussion

MCPyV’s tumorigenic effect is most lethal in immunosuppressed patients [14, 15], highlighting the critical role of host immunity in controlling virus’s oncogenic potential. However, the mechanisms through which MCPyV interacts with host innate immunity to sustain an asymptomatic persistent infection, and how a failure to control of MCPyV infection leads to MCC tumorigenesis, remain significant yet poorly understood issues.

In this study, we uncovered a mechanism through which MCPyV modulates the host STING-TBK1 signaling pathway, achieving tightly controlled viral infection. We discovered that among all the MCPyV-encoded proteins, only ALTO can stimulate STING downstream IFN production (Fig 1 and S1 Fig). The most notable effect of ALTO expression is the initiation of TBK1 autophosphorylation, ultimately leading to the stabilization of STING and the activation of downstream components like IRF3 and NF-κB (Fig 2, S2 and S3). Using several complementary approaches, including Co-IP, NanoBiT, and BiFC studies, we found that ALTO interacts with both STING and TBK1 (Fig 3, S4 and S6). Molecular analysis showed that ALTO contains multiple PXXP motifs that are recognized by the SH3 domain of the Src tyrosine kinase (Fig 4). We further demonstrated that ALTO likely recruits Src into the TBK1 complex through its PXXP motifs to enhance TBK1 phosphorylation (Fig 4 and S10 Fig), thereby activating STING pathway and its downstream IFN signaling. Interestingly, unlike STING activation by diABZI, which leads to its degradation, MCPyV infection stabilizes STING protein (Fig 5), hinting that ALTO protein expressed in MCPyV-infected cells may play a role in this stabilization. Indeed, our analysis of MCPyV-infected HDFs showed a physiological significance of ALTO’s activity during viral infection. Loss- and gain-of-function studies in MCPyV-infected HDFs indicated that ALTO suppresses MCPyV infective activity and that TBK1 is crucial in mediating ALTO’s function in controlling MCPyV infection (Fig 6-8, and S13 Fig). Thus, our findings suggest that the interaction between ALTO and the STING- TBK1 pathway could be a strategy by which MCPyV circumvents host defenses, limiting its infection activity to maintain a balanced coexistence with the host immune system, enabling persistent infection (Fig 8C).

The cGAS-STING pathway is a crucial component of the innate immune system, detecting viral DNA in the cytoplasm and initiating a signaling cascade that ultimately leads to IFN production and an antiviral immune response. This pathway often becomes a prime target for degradation or inhibition by viral proteins. In the literature, the most extensively studied mechanisms focus on how viral effectors target and suppress innate immune factors, thereby promoting a proviral state conducive to either rapid expansion during acute infections or latent/chronic persistence within the host cell [47, 48]. However, in this paper, we describe a unique strategy in which a viral protein stimulates host innate immune factors, activating an antiviral state to regulate viral propagation. We have shown that ALTO effectively stimulates TBK1 autophosphorylation, thereby priming the STING signaling pathway. This mechanism ensures that the negative regulation of MCPyV infectivity is initiated only after the virus has achieved sufficient replication within the host cell nucleus, allowing for adequate cytoplasmic expression of ALTO to activate the STING-TBK1 pathway. This concept is reinforced by our observations that STING downstream cytokines and ISGs are induced only after the completion of viral replication and transcription [33], enabling the ALTO protein to activate the STING signaling pathway. Further supporting this, we found that the ability of ALTO to trigger TBK1 autophosphorylation is enhanced by the STING agonist diABZI, suggesting that ALTO’s activity is likely induced by active MCPyV replication. Even minimal amounts of ALTO can rapidly stimulate the STING-TBK1 pathway (Fig 1, S2 and S3 Fig), implying that intracellular levels of ALTO protein may act as a sort of “emergency brake.” As the virus undergoes replication and transcription, the initial expression of ALTO might be a low-probability event. However, as viral replication progresses, increasing amounts of ALTO accumulate in the cytosol, priming the STING-TBK1 pathway for sustained activation. Analysis of single-cell ALTO phenotypes during MCPyV infection revealed that MCPyV-infected cell population displayed a significantly variable level of ALTO protein, likely due do the timing of viral infection in specific cells. We also found that ALTO represses MCPyV infection at the single cell level. Specifically, ALTO-positive HDFs often displayed significantly fewer viral proteins and replicated genomes compared to ALTO-negative cells (Fig 7 and S13 Fig). This result further supports a negative-feedback mechanism mediated by ALTO, allowing each cell to respond to its specific viral load while exerting a direct ’autocrine’ effect to control viral propagation. Despite MCPyV’s almost Spartan genome, which contains only a handful of distinct genes, the overprinting of its primary antigen has enabled the virus to produce a regulatory protein that acts not directly on the virus itself but on its cellular environment to prevent over-replication. Thus, this study unveils a unique strategy for the virus to maintain persistent infection within its host.

We have also demonstrated that Src kinase plays a critical role in ALTO- induced TBK1 phosphorylation and downstream signaling (Fig 4). In the human skin, activated Src is involved in regulating cell adhesion, migration, and proliferation in processes such as wound healing and skin regeneration [49, 50]. Given the specific cellular environment within the human skin and the activity of Src in these cells, it’s plausible that there could be a cell type-specific mechanism to modulate the ALTO-induced antiviral response. Indeed, recent findings from the Chang and Moore laboratory underscore the close relationship between Src, the SFKs, and ALTO [8], highlighting the importance of studying this relationship in MCPyV biology. Future studies will explore how skin damage induced by UV/ionizing radiation and the subsequent wound healing processes might alter Src function and impact MCPyV infection in the physiological skin environment.

Our studies suggest that in normal cells, STING signaling is tightly regulated to restrict MCPyV infection, thus establishing a balance between the virus and host that supports persistent infection. Previously, we also found that STING is silenced in MCPyV(+) MCC tumors [34], indicating that the STING-mediated innate immune response might function as an intrinsic barrier to MCPyV-driven tumorigenesis. It is conceivable that a compromised STING function could lead to unrestrained MCPyV replication and integration, thereby promoting the development of MCPyV-driven MCC. The work described herein lays the groundwork for investigating how the dysregulation of STING signaling, induced by aging or compromised host immunity, might alter the interaction between MCPyV and its host cell, potentially facilitating virus-induced malignancy. These studies could reveal novel strategies for preventing and treating devastating MCC cancers.

## Materials and Methods

### Cell culture

HDFs were isolated from neonatal foreskins as previously described [28, 30] and were maintained in DMEM (Invitrogen) supplemented with 10% FBS (Cytiva), 1x nonessential amino acids (Gibco), and 1x GlutaMAX (Gibco).

HEK293 cells were maintained in DMEM (Invitrogen) supplemented with 10% FBS (Cytiva). U2OS cells were maintained in McCoy 5A (Invitrogen) supplemented with 10% FBS (Cytiva). All cells were maintained and incubated at 37°C in 5% CO2.

### Chemicals and reagents

Saracatanib, Dasatanib, Bosutinib, BX-795, and GSK8612 (SelleckChem) were dissolved in DMSO to a stock concentration of 10mM. diABZI (SelleckChem) was dissolved in DMSO to a stock concentration of 5mM. diABZI was used at a final concentration of 1 μM in all experiments. Doxycycline (Clontech) was dissolved in water to a stock concentration of 0.5mg/mL. Dox was used at a final concentration of 0.5 μg/ml in all experiments unless otherwise indicated in the figure legend. Stocks of diABZI and Doxycycline were stored as small aliquots at -20°C. All other stock solutions of chemicals were stored in small aliquots at −80°C.

### Recombinant plasmid construction

The pTRIPZ-STING R284S construct was generated in our previous studies [36]. The viral gene constructs used in this study were sourced as follows: pGwf and phGf were from Addgene; pALTOw, pMtw (ST), pWm (VP1), pMono-Zeo, pMono-ADL (LT) and ph2m (VP2) were from the Christopher Buck Laboratory; and the pcDNA4c-57kT construct was generated by first fusing the full 57kT ORF using two-step PCR from pcDNA4c-LT1-817 [51] and then cloning it into the pcDNA4c backbone using the KpnI and XhoI restriction sites.

For making HDF-inALTO stable cells, the ALTO coding sequence was PCR-amplified from pALTOw (Christopher Buck Laboratory) and then cloned using AgeI and MluI sites to replace the RFP coding sequence in the pTRIPZ vector (Open Biosystems). The resulting construct is named pTRIPZ-ALTO.

To generate the ALTOnull MCPyV genome, the region surrounding the ALTO coding sequence in pR17b (WT MCPyV, Christopher Buck Laboratory), between the restriction sites SacI and HapI, was amplified by two-step PCR to mutate the ALTO ATG start codon to ACG, without altering the Large T sequence or reading frame. This altered sequence was then subcloned back into the viral plasmid using those same restriction sites.

NanoBiT plasmids pBiT1.1-N[TK/LgBiT], pBiT1.1-C[TK/LgBiT], pBiT2.1-N[TK/SmBiT], and pBiT2.1-C[TK/SmBiT] were obtained from Promega. ALTO, STING, and TBK1 sequences were PCR amplified from the pALTOw plasmid (Christopher Buck Laboratory) or normal HDF cDNA samples and inserted into each NanoBiT vector using the XhoI and NheI (for ALTO and STING) or XhoI and BglII sites (for TBK1).

BiFC constructs were cloned using the same strategy as described in our previous study [52, 53]. Coding sequences for ALTO (PCR-amplified from pALTOw), TBK1 and STING (both PCR-amplified from the NanoBiT expression plasmids), were cloned using XhoI and NotI sites in the BiFC vectors to generate the in-frame fusion of the ALTO/TBK1/STING molecules with N- or C-terminal Venus N or Venus C.

### Preparation of HDF inducible stable cells

Inducible RFP or ALTO stable cells were generated from early passage primary HDFs (≤ 2 passages from tissue) as previously described [33, 34, 54]. Tetracycline-screened FBS (Cytiva) was used to supplement the cell culture media as described above.

### MCPyV infection

ALTOnull MCPyV was prepared using the WT MCPyV method described previously [28, 30]. Normal HDFs or HDF-inALTO stable cells were infected with WT or ALTOnull MCPyV using the method described previously [28, 30, 54]. For virus infection, unless specifically noted in the figure legends, 10^8^ viral genome equivalents of MCPyV virions were used in each 96-well.

To analyze MCPyV replication, harvested cell pellets were lysed in QuickExtract DNA Extraction Solution (Biosearch Technologies) according to the manufacturer’s instructions. The extracted DNA samples were then subjected to real-time qPCR using a QuantStudio 3 Real-Time PCR System (Applied Biosystems) with SYBR Green Master Mix (Applied Biosystems). The MCPyV genomic DNA levels were measured using NCRR-targeted primers and normalized to the level of genomic GAPDH. Both NCRR and genomic GAPDH primer sequences can be found in [33].

### Cell viability assay

HDF-inALTO cells were maintained as described above in Tet-free media. A total of 2x10^4^ cells were seeded in wells of a 96-well dish and allowed to adhere for 8 hours at 37°C in 5% CO2. Standard media was changed to media containing 0.5 μg/ml Dox and the tested concentration of each drug. After incubating at 37°C in 5% CO2 for 16 hours, cell viability assays were performed using the Cell Titer Glo 3D kit (Promega) according to manufacturer instructions. Luminescence was measured using a Luminoskan Ascent microplate reader (Thermo). Relative viability was calculated by comparing average luminescence readings between drug-treated and DMSO vehicle control-treated cells.

### NanoBiT protein-protein interaction assay

HEK293 cells were seeded at 3x10^4^ cells in 100µL per 96-well approximately 20 hours prior to transfection. 100ng total DNA of the examined NanoBiT fusion protein plasmids (50ng LgBiT + 50ng SmBiT) were diluted in Opti-MEM I (Gibco) and combined with diluted Lipofectamine 2000 (Invitrogen) at a 2.2:1 ratio of Lipofectamine to DNA before being added to cells. Transfected cells were incubated at 37°C in 5% CO2 for 20-24 hours. Approximately 30 minutes prior to desired reading time, cell media was aspirated and replaced with 100µL Opti-MEM I (Gibco). NanoGlo reagent was freshly diluted to a 5X stock in Live Cell Substrate according to manufacturer direction, then added to all wells. Plates were returned to the incubator for 15 minutes, and luminescence was measured using a Luminoskan Ascent microplate reader (Thermo). Per manufacturer direction, a specific protein-protein interaction was assessed to be present when the average luminescence reading of the interacting pair was tenfold higher than the average from the LgBiT-tagged construct co-transfected with the SmBiT negative control construct.

### BiFC assay

1.5x10^5^ U2OS cells were seeded in a 6-well plate atop 1-2 coverslips and allowed to adhere overnight at 37°C in 5% CO2. Subsequently, 1 μg of each construct of the BiFC pair for each well was diluted in Opti-MEM I (Gibco). 6.5 μL FuGENE HD (Promega) was added to this mixture and allowed to incubate for 15 minutes at RT prior to addition to cells. BiFC constructs were incubated at 37°C in 5% CO2 for 24-48 hours prior to cellular fixation and IF staining.

### Immunofluorescent staining

Cells were cultured on coverslips in standard conditions and fixed with 3% paraformaldehyde in PBS for 10-20 minutes. Staining was performed as previously described [33]. Primary antibodies used in this study are as follows: anti-ALTO (1:10,000, Denise Galloway Lab), anti-MCPyV LT (1:250, sc-136172, Santa Cruz Biotechnology), anti-STING (1:200, 19851-1-AP, Proteintech), anti- Mx1 (1:1,000, 37849S, Cell Signaling Technology), and anti-FLAG (1:40,000, F3165, Sigma Aldrich). Secondary antibodies used in this study are as follows: Alexa Fluor 594 goat anti-mouse IgG (1:500, A11032, Invitrogen), Alexa Fluor 488 goat anti-rabbit IgG (1:500, A11034, Invitrogen), Alexa Fluor 594 goat anti-rabbit IgG (1:500, A11012, Invitrogen), and Alexa Fluor 488 goat anti-mouse IgG (1:500, A11029, Invitrogen).

All IF images were captured using an inverted fluorescence microscope (IX81; Olympus) as described previously [33]. The scale bars were added using ImageJ software.

### Immunofluorescent-fluorescence in situ hybridization (ImmunoFISH)

HDFs were fixed with 3.7% formaldehyde in PBS for 10 minutes. IF staining was performed using ALTO antibody with 1:3,000 dilution as previously described [33]. After IF staining, the cells were fixed with 3.7% formaldehyde in PBS for 10 minutes again and subjected to FISH staining using buffers from Biosearch Technologies as previously described [33]. After the final wash step, cells were mounted on coverslips and observed using an inverted fluorescence microscope (IX81; Olympus).

The FISH signals were quantified at the single cell level using ImageJ. Photos from each of the channels of interest were stacked. Using the DAPI channel, the nuclear area of a single ALTO- or FISH-positive cell was selected. Both ALTO- and FISH-positive cells were considered infection-positive. The area selection was then applied to the FISH channel and the integrated density (the sum of the signal intensity in the selected area) was measured for that cell. More than 40 cells were analyzed for each group of samples to achieve a statistically meaningful sample size.

### Structural prediction

The canonical ALTO sequence, with a length of 250 amino acids, was utilized to generate the predicted structure. The tool AlphaFold 2 processed this sequence on the public Google Colab (ColabFold v1.5.5: AlphaFold2 using MMseqs2) using default settings, without any constraints, templates, or multimerization options.

### Western blotting

Whole cell lysate preparation and Western blot analyses were performed as previously described [33]. Primary antibodies utilized in this study are as follows: anti-GAPDH (1:5,000, 2118S, Cell Signaling Technology), anti-ALTO (1:10,000, Denise Galloway Laboratory), anti-STING (1:1000, 13647S, Cell Signaling Technology), anti-STING p-S366 (1:1000, 19781S, Cell Signaling Technology), anti-TBK1 (1:1000, 3504S, Cell Signaling Technology), anti-TBK1 p-S172 (1:1000, 5483S, Cell Signaling Technology), anti-P65 (1:1000, 8242S, Cell Signaling Technology), anti-P65 p-S536 (1:1000, 3033S, Cell Signaling Technology), anti-IRF3 (1:500, SC-33641, Santa Cruz Biotechnology), anti-IRF3 p-S386 (1:1000, 37829S, Cell Signaling Technology), anti-RFP (1:2,000, AB233, Evrogen), rabbit anti-Src antibody (1:1000, 2109S, Cell Signaling Technology), mouse anti-Src antibody (1:1000, 2110S, Cell Signaling Technology), anti-LgBiT (1:500, N7100, Promega), anti-MCPyV LT (1:500, sc-136172, Santa Cruz Biotechnology), and anti-MCPyV VP1 (1:2,500, Christopher Buck Laboratory).

The secondary antibodies utilized in this study are HRP-linked anti-rabbit IgG (1:2000, 7074S, Cell Signaling Technology), and HRP-linked anti-mouse IgG (1:2000, 7076S, Cell Signaling Technology). Western blots were developed with the SuperSignal West Pico PLUS Chemiluminescent Substrate Kit (Thermo Fisher Scientific) and imaged using an Amersham Imager 600 or 680 (GE Healthcare).

Quantification of signals in Western blots was performed using ImageJ. Briefly, each lane was defined using a tall, narrow rectangle. A profile plot for each band within the lane was generated using the Plot Lanes function. The peak area for each lane was closed with a straight line and measured using the wand tool. Relative abundance was calculated by setting the value of a control band as 1.

### RT-qPCR

Total RNA was isolated from cell samples using the NucleoSpin RNA XS Kit (Macherey-Nagel) following the manufacturer’s instructions. Reverse transcription and Realtime qPCR were performed as previously described [54].

### Statistical analyses

Unpaired, two-tailed t-tests were performed using the t.test function in Microsoft Excel. The significance thresholds were set at *P value < 0.05, **P < 0.01, ***P < 0.001.

## Acknowledgments

The authors would like to express their gratitude to Christopher Buck (NCI) for providing pR17b and the plasmids encoding MCPyV genes, and to Deyuan Xu and Qixun Sun for technical support in preparing the BiFC constructs. Our appreciation goes to the members of our laboratories for their valuable discussions.

## Supporting information

**S1 Fig.**
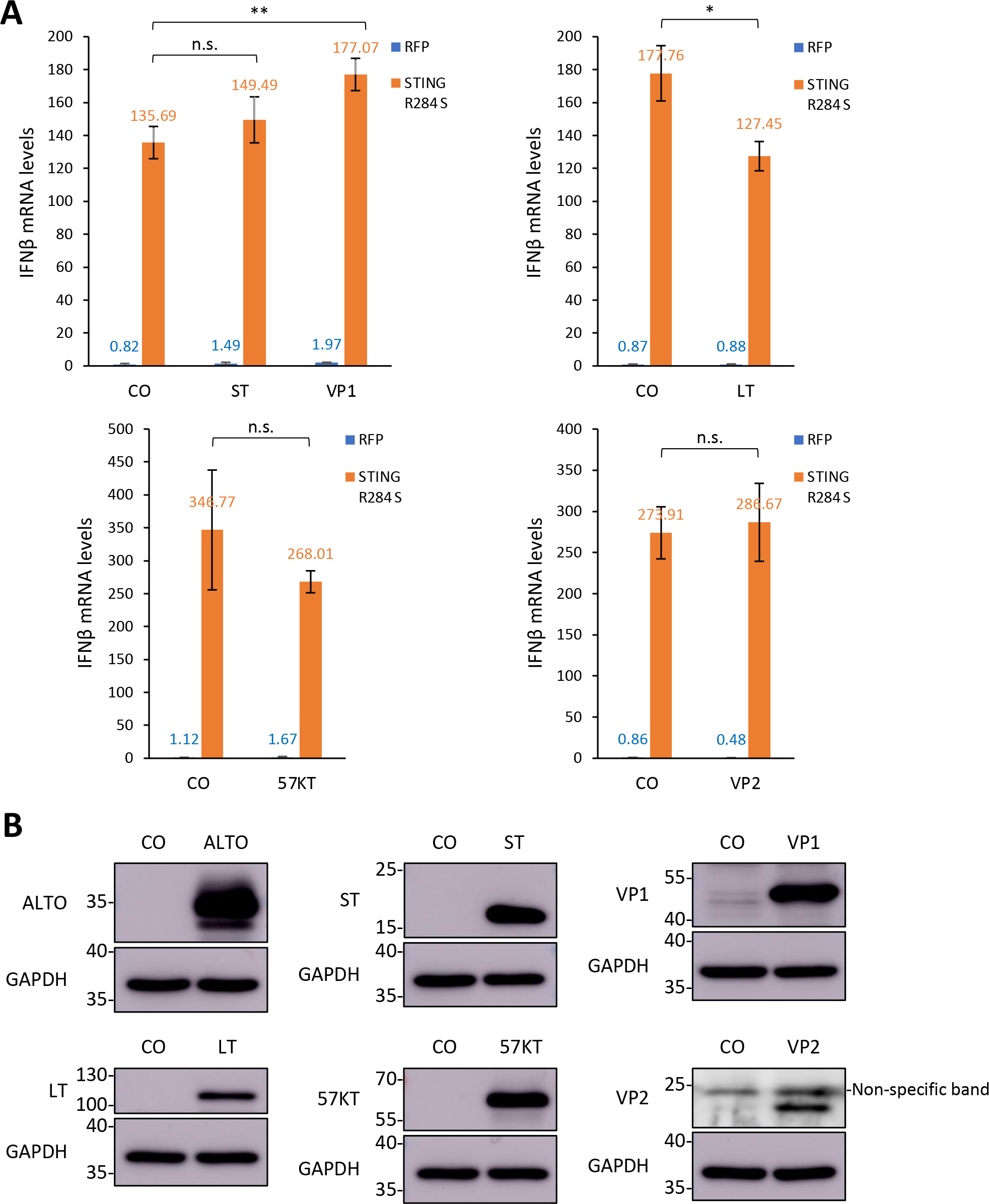
ALTO is the only MCPyV protein that could stimulate STING signaling. A. HEK293 cells were transfected with plasmids carrying indicated viral proteins (or empty vector controls) and STING R284S (or RFP control). At 2 days post-transfection, IFNβ mRNA levels were measured using RT-qPCR. One of the values for transfection with vector and RFP was set as 1. Error bars indicate standard deviation from three independent samples. **p<0.01; *p<0.05; n.s. = not significant. **B**. HEK293 cells were transfected with plasmids encoding the indicated viral proteins or their respective empty vector controls. Whole cell lysates were then blotted with the indicated antibodies to confirm the expression of the viral proteins.

**S2 Fig.**
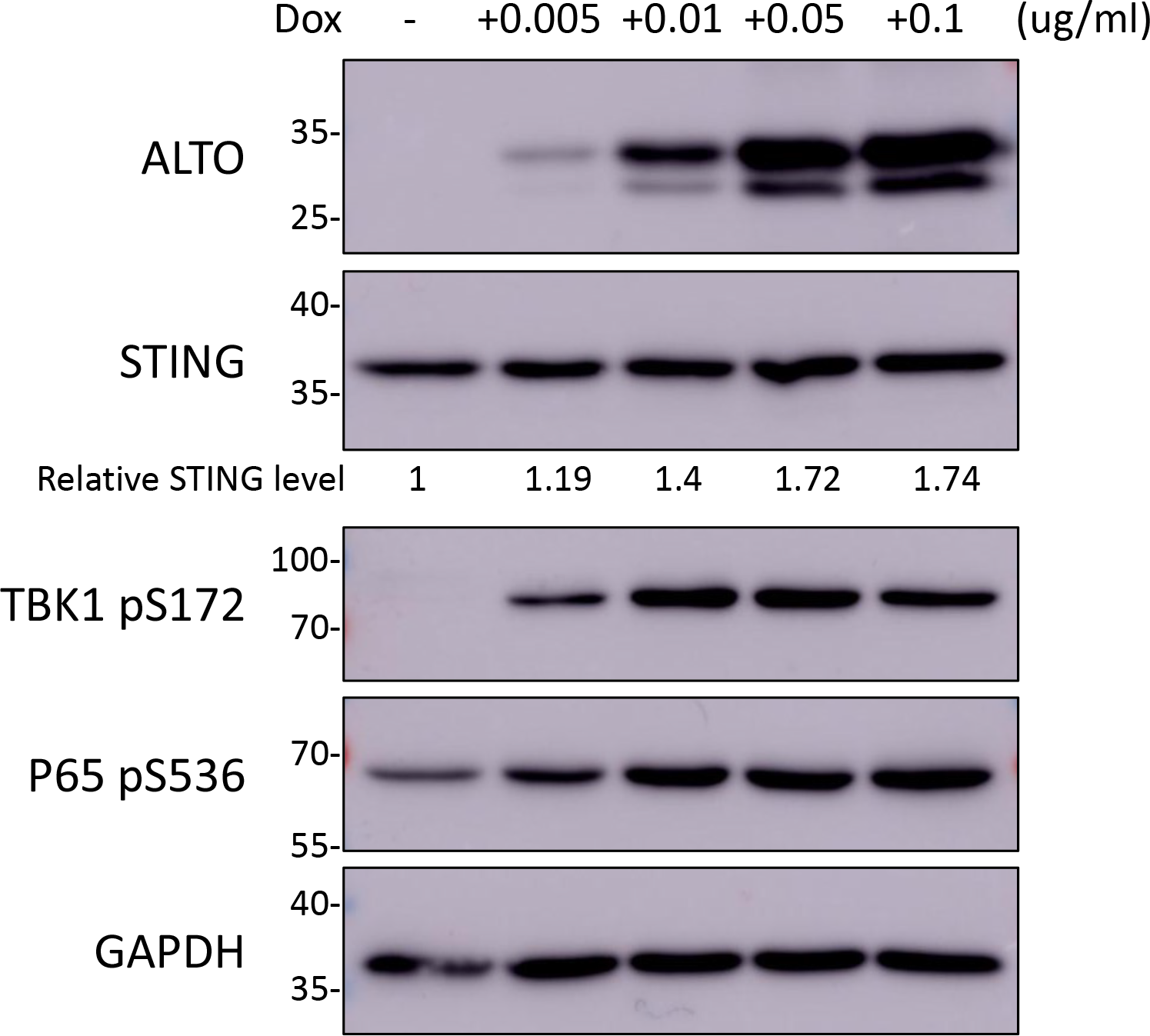
A trace amount of ALTO can activate TBK1^S172^ autophosphorylation. HDF-inALTO cells were mock-treated or induced with the indicated doses of Dox for 42 hours. Whole cell lysates were resolved by SDS/PAGE and immunoblotted with the indicated antibodies. Relative abundance of STING was calculated in ImageJ by setting the value of the band in the uninduced cells to 1.

**S3 Fig.**
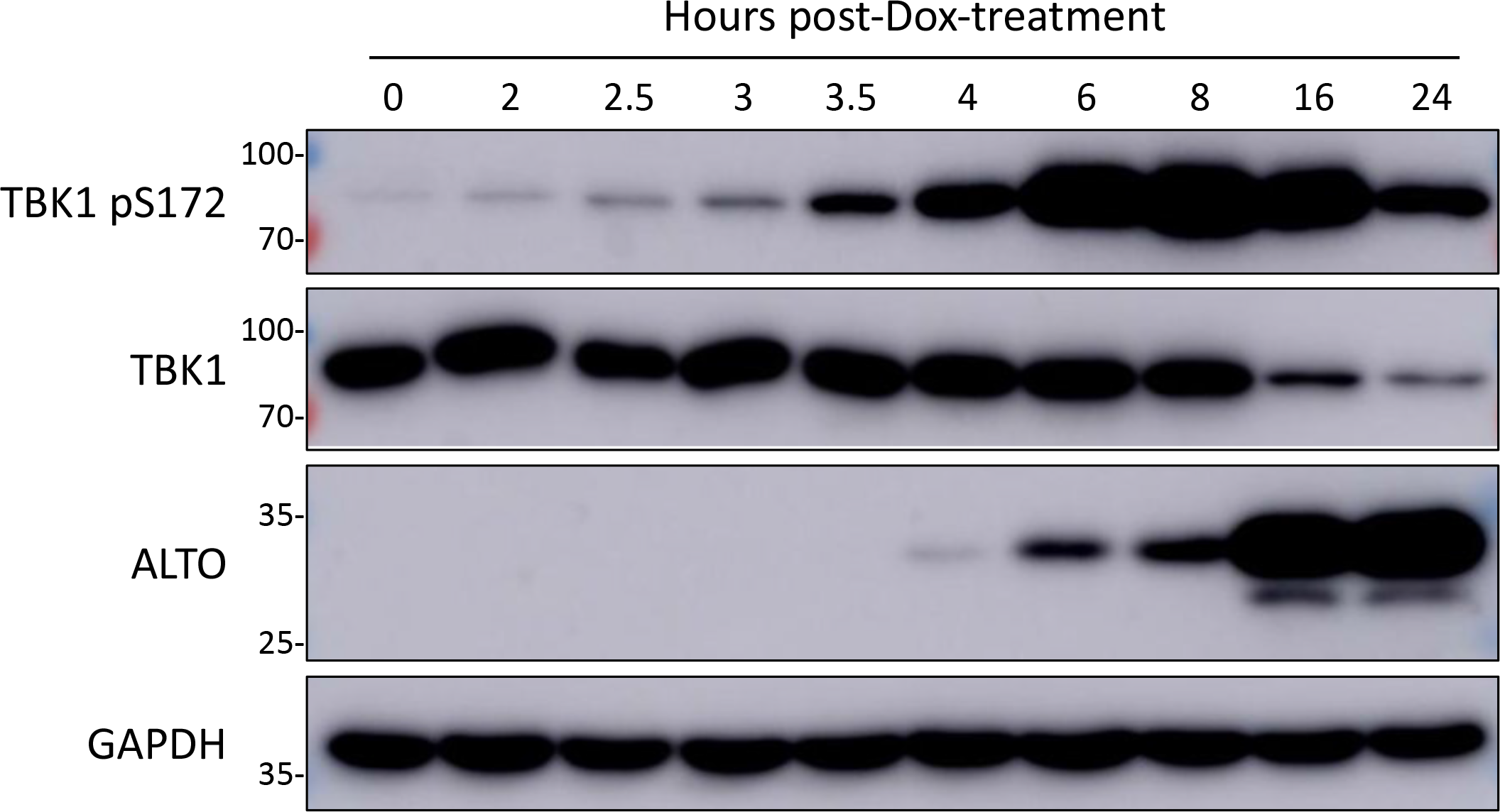
ALTO rapidly induces TBK1^S172^ autophosphorylation. HDF-inALTO cells were mock-treated or induced with Dox for the indicated times. Whole cell lysates were resolved by SDS/PAGE and immunoblotted with the indicated antibodies.

**S4 Fig.**
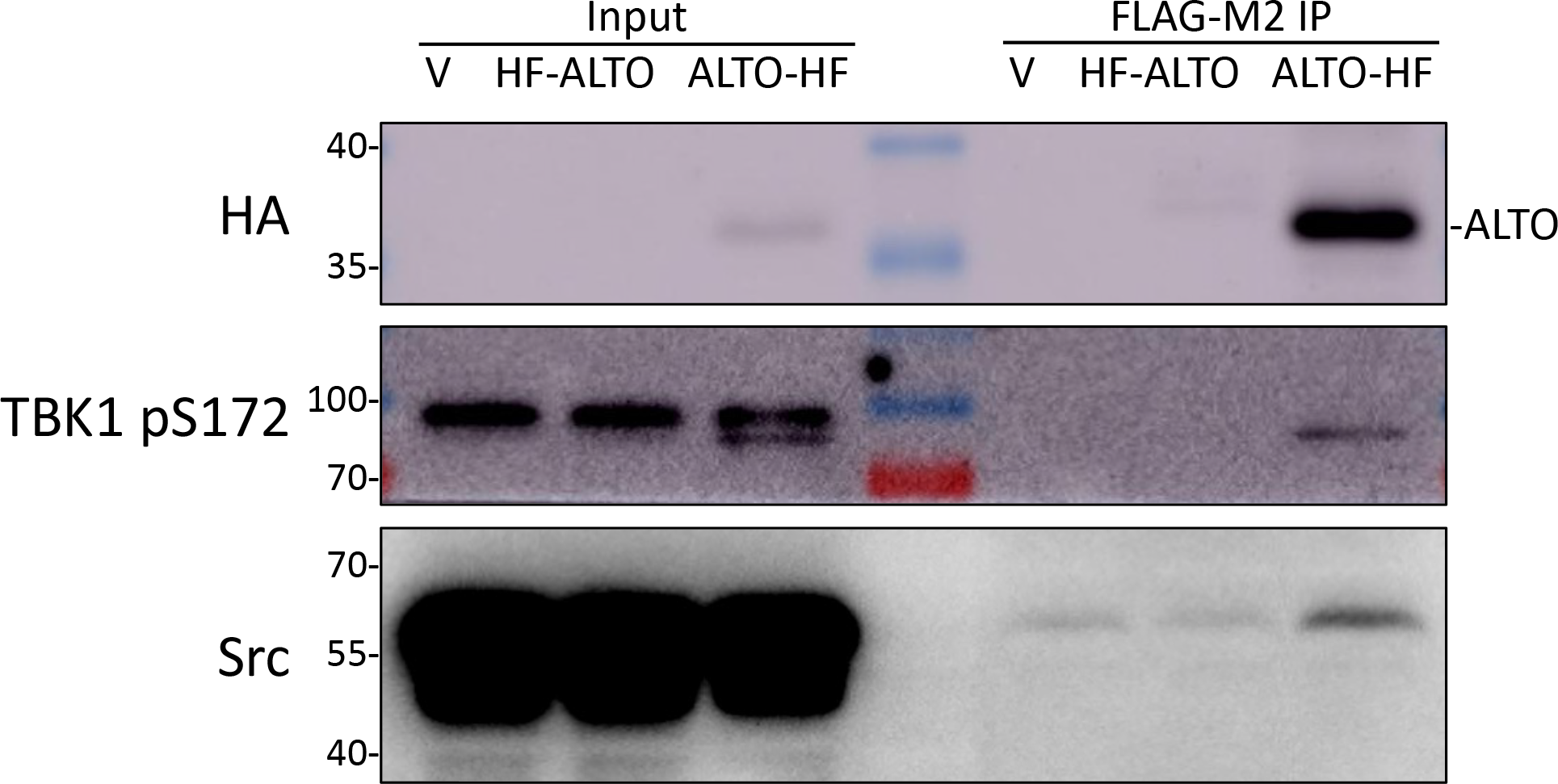
ALTO interacts with both S172 phosphorylated TBK1 and Src. HEK293 cells were transfected with empty vector control or vector containing N- or C-terminal HA- and FLAG-tagged ALTO. At 40-hour post-transfection, whole cell lysates were incubated with Anti-FLAG M2 affinity gel. Immunoprecipitants were resolved by SDS/PAGE and immunoblotted with the indicated antibodies.

**S5 Fig.**
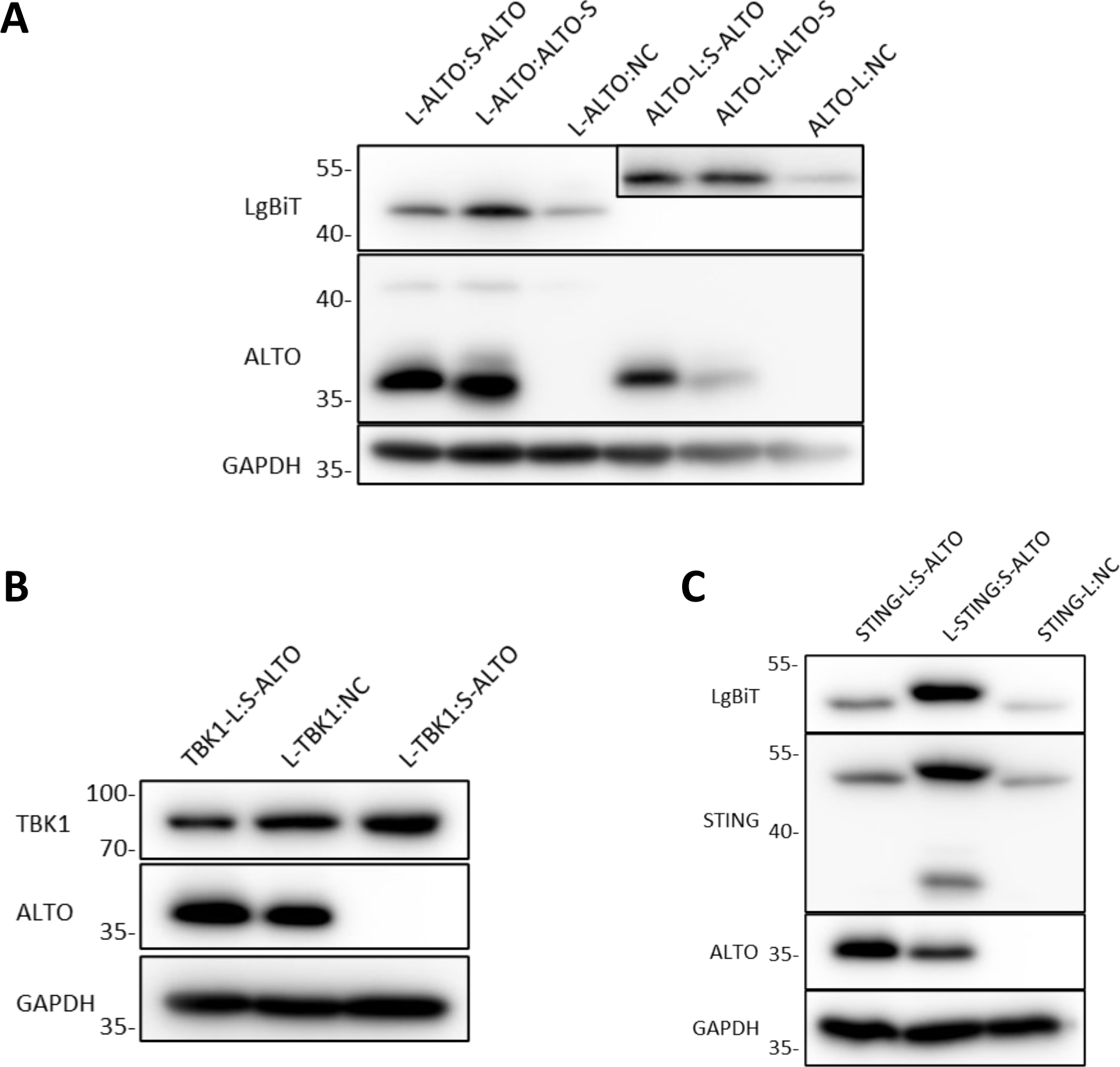
The NanoBiT fusion proteins were co-expressed in cells. A. HEK293 cells were transfected with pairs of constructs carrying NanoBiT- (Large BiT or SmBiT) fused ALTO as indicated. At 24 hours post-transfection, whole cell lysates were collected and resolved by SDS/PAGE and immunoblotted with the indicated antibodies. Inset: To enhance visibility of the faint ALTO-LgBiT bands in the last three lanes, the LgBiT-blotted membrane was re-imaged with the LgBiT- ALTO bands in the first three lanes covered and the exposure time increased. **B.** HEK293 cells were transfected with pairs of constructs carrying NanoBiT- (Large BiT or SmBiT) fused ALTO and TBK1 as indicated. At 20 hours post-transfection, whole cell lysates were collected and resolved by SDS/PAGE and immunoblotted with the indicated antibodies. **C.** HEK293 cells were transfected with pairs of constructs carrying NanoBiT- (Large BiT or SmBiT) fused ALTO and STING as indicated. At 24 hours post-transfection, whole cell lysates were collected and resolved by SDS/PAGE and immunoblotted with the indicated antibodies.

**S6 Fig.**
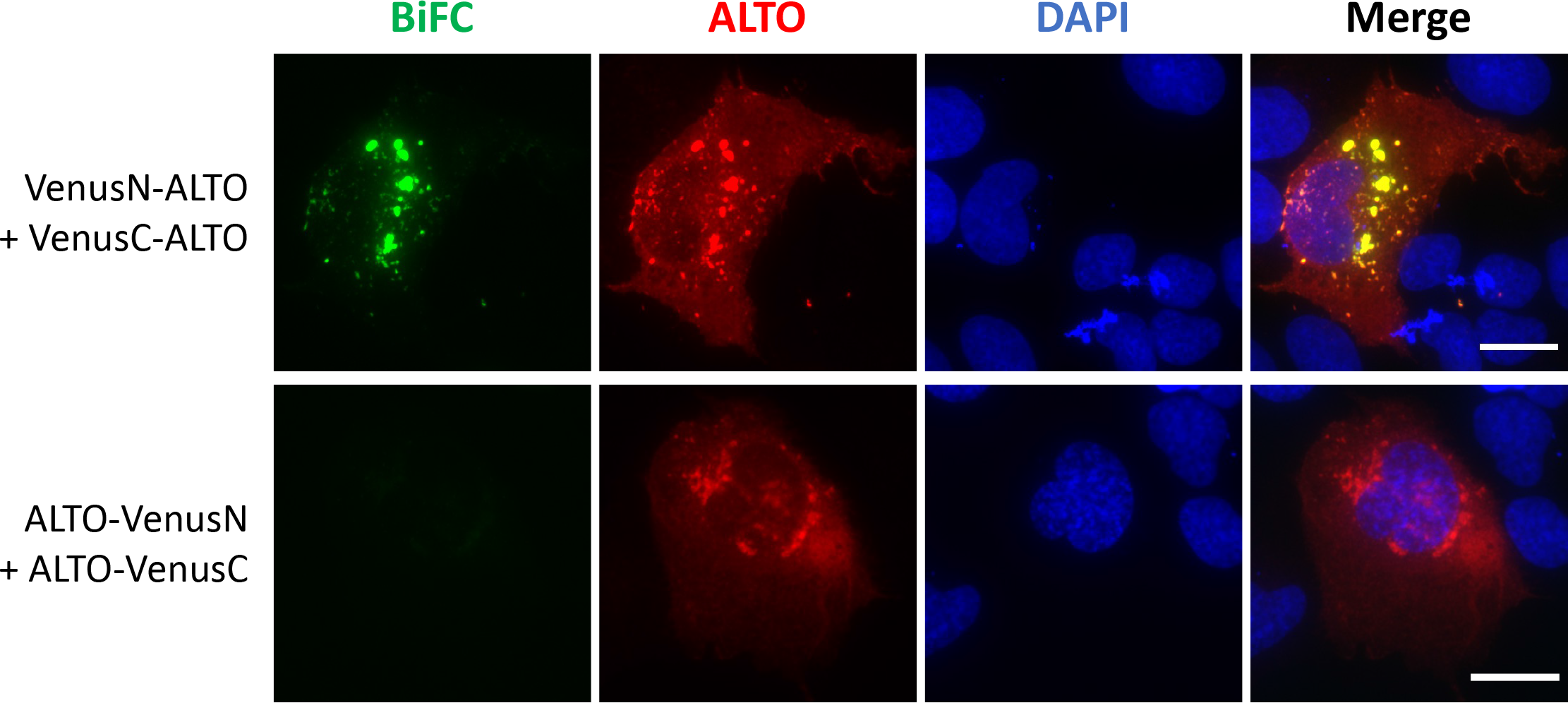
BiFC detection of ALTO-ALTO interaction. U2OS cells were transfected with pairs of plasmids carrying Venus (N-terminal half or C-terminal half) fused ALTO constructs. At 48-hour post-transfection, cells were fixed and immunostained for ALTO, and counterstained with DAPI. Scale bar, 20 μm.

**S7 Fig.**
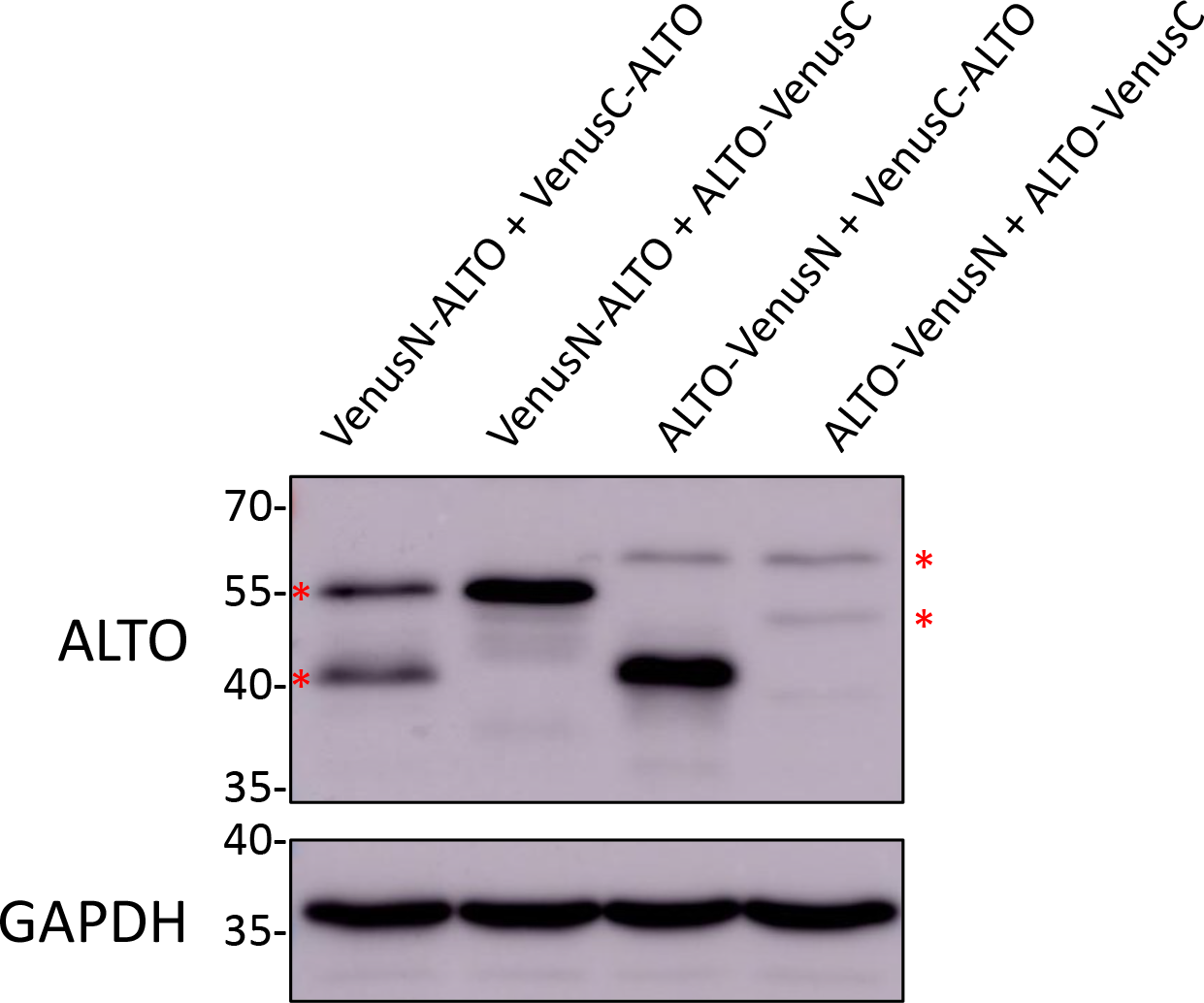
ALTO-Venus fusion proteins were efficiently expressed in cells. U2OS cells were transfected with pairs of plasmids carrying Venus (N-terminal half or C-terminal half) fused ALTO constructs as indicated. At 48-hour post-transfection, whole cell lysates were resolved by SDS/PAGE and immunoblotted with the indicated antibodies. Red asterisks indicate bands that reflect the position of full-length ALTO protein, which was variably shifted due to tagging.

**S8 Fig.**
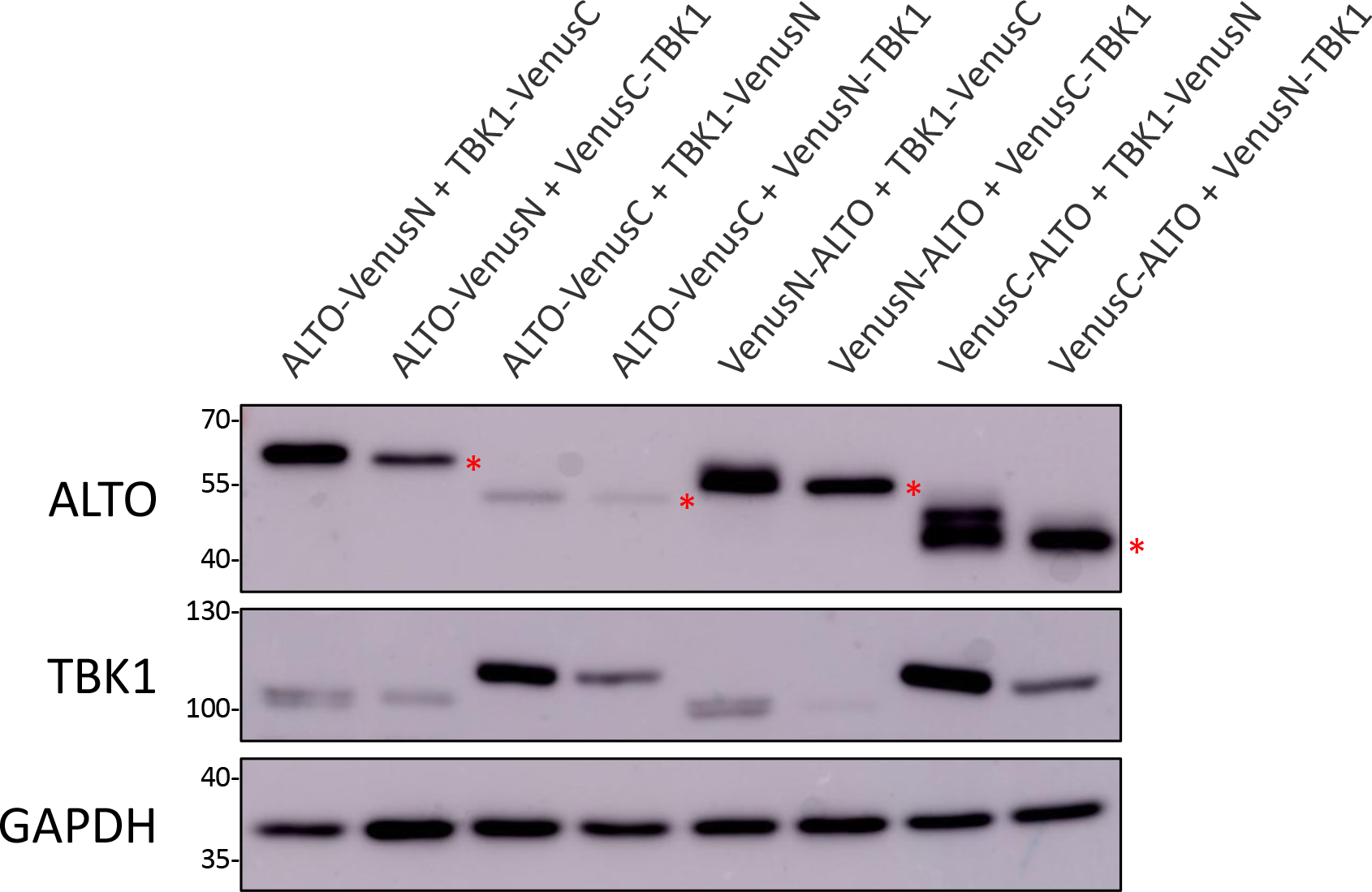
Venus-tagged ALTO and TBK1 were efficiently expressed in cells. U2OS cells were transfected with pairs of plasmids carrying Venus (N-terminal half or C-terminal half) fused ALTO and TBK1. At 24-hour post-transfection, whole cell lysates were resolved by SDS/PAGE and immunoblotted with the indicated antibodies. Red asterisks indicate bands that reflect the position of full-length ALTO protein, which was variably shifted due to tagging.

**S9 Fig.**
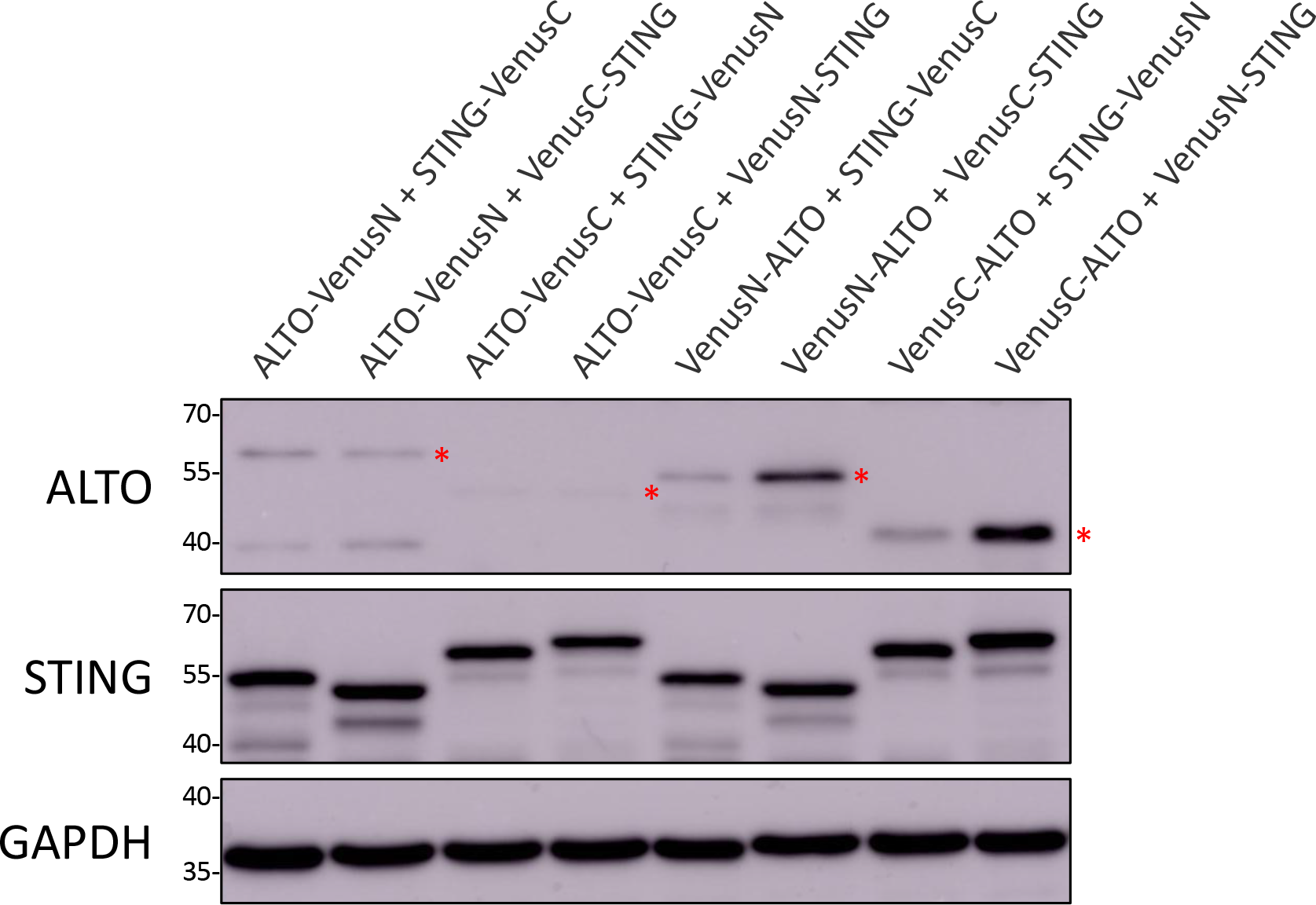
Venus-tagged ALTO and STING were efficiently expressed in cells. U2OS cells were transfected with pairs of plasmids carrying Venus (N-terminal half or C-terminal half)-fused ALTO and STING. At 24-hour post-transfection, whole cell lysates were resolved by SDS/PAGE and immunoblotted with the indicated antibodies. Red asterisks indicate bands that reflect the position of full-length ALTO protein, which was variably shifted due to tagging.

**S10 Fig.**
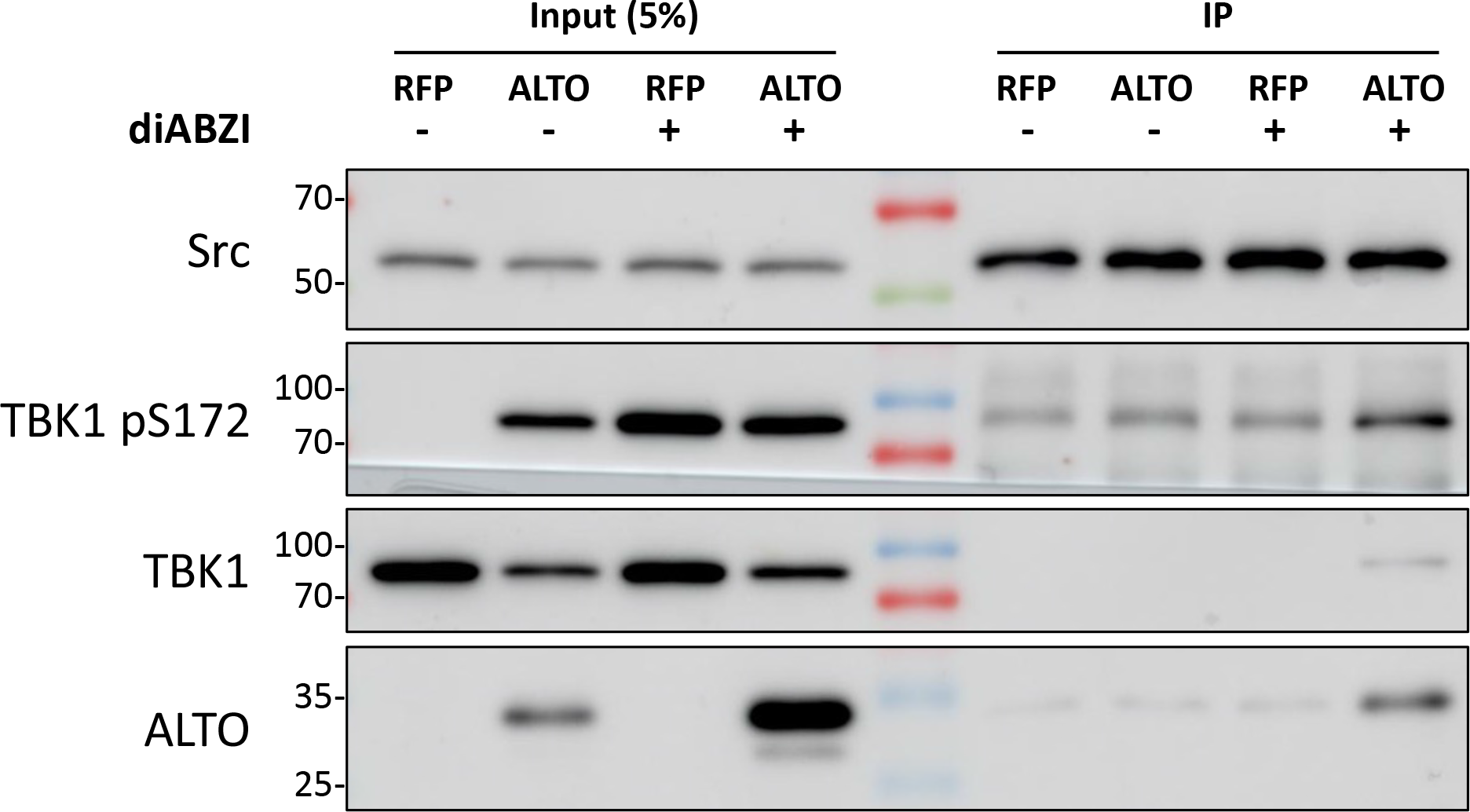
ALTO enhances Src and TBK1 association. Normal HDF-inALTO or - inRFP cells were mock-treated or induced with Dox for 10.5 hours, then stimulated with diABZI (or DMSO control) for 1.5 hours. Whole cell lysates were incubated with rabbit anti-Src antibody, then immunoprecipitated with Protein G agarose. Immunoprecipitants were resolved by SDS/PAGE and immunoblotted with the indicated antibodies.

**S11 Fig.**
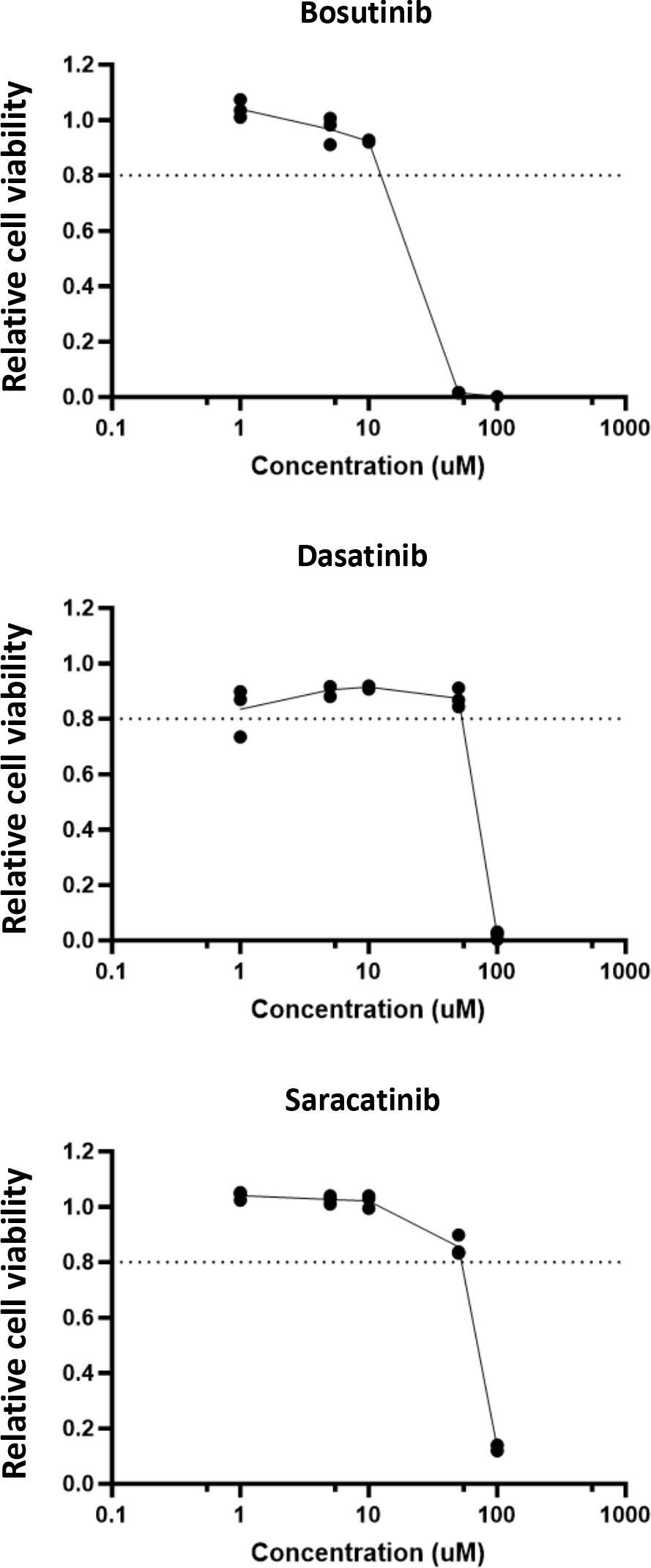
Src/SFK inhibitors are well-tolerated by HDFs. Normal HDF-inALTO cells were co-treated with Dox and the indicated concentrations of Src/SFK inhibitor for 16 hours. Cell viability was assessed via luminescence using Cell Titer Glo 3D; relative viability was calculated by comparing drug-treated cells to DMSO-treatment. Points indicate replicate wells.

**S12 Fig.**
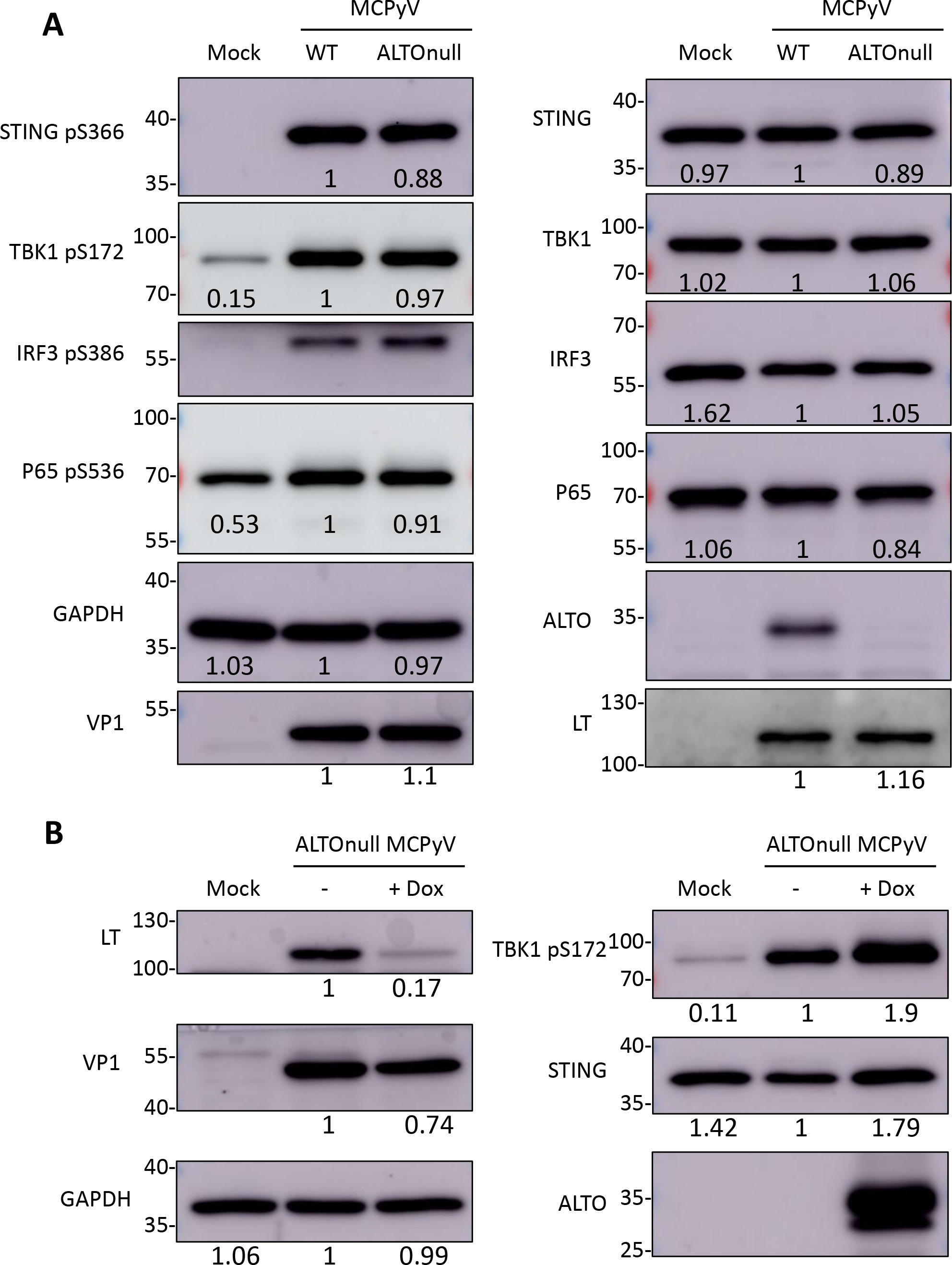
Activation of STING and downstream effectors in HDFs infected with WT and ALTOnull MCPyV. A. Normal HDFs were mock-infected or infected with WT or ALTOnull MCPyV. Whole cell lysates were collected on day 5 post-infection, resolved by SDS/PAGE, and immunoblotted with the indicated antibodies. Quantifications were performed in ImageJ by setting the value of the WT infection to 1. **B.** HDF-inALTO cells were infected with ALTOnull MCPyV. Cells were either mock-treated or induced with Dox on days 2 and 4 post-infection. Whole cell lysates were collected on day 5 post-infection, resolved by SDS/PAGE, and immunoblotted with the indicated antibodies. Quantifications were performed in ImageJ by setting the value of the no-Dox infection control to 1.

**S13 Fig.**
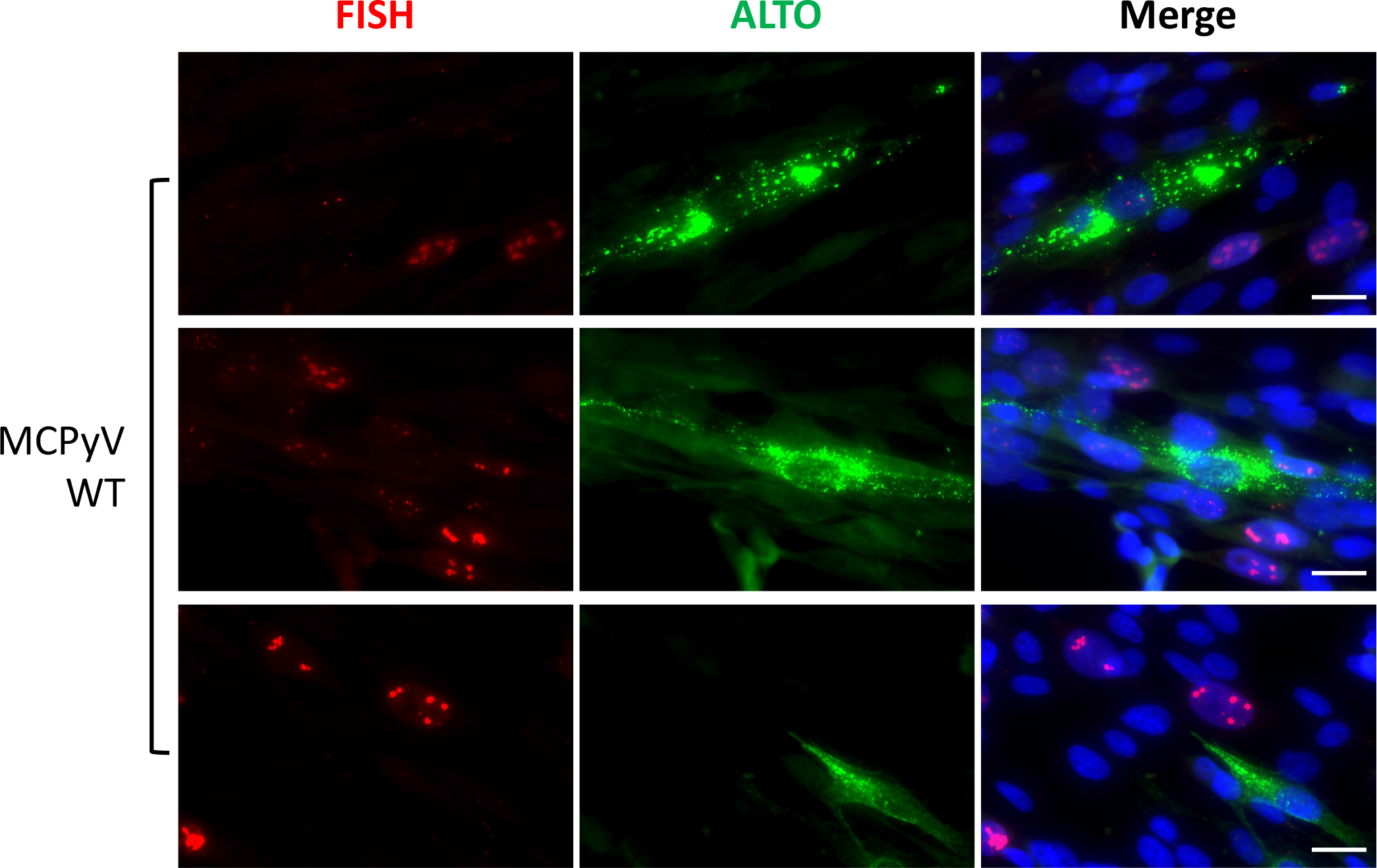
ALTO-positive cells show reduced viral replication. Normal HDFs were mock-infected or infected with 6x10^8^ copies of WT or ALTOnull MCPyV per 48-well. On day 5 post-infection, cells were fixed and immunostained for ALTO, then subjected to FISH using MCPyV probe, and finally counterstained with DAPI. Scale bar, 20 μm.

